# Adenosine triggers early astrocyte reactivity that provokes microglial activation and drives the pathogenesis of sepsis-associated encephalopathy

**DOI:** 10.1101/2023.10.30.563169

**Authors:** Qilin Guo, Davide Gobbo, Na Zhao, Qing Liu, Li-Pao Fang, Tanja M. Gampfer, Markus R. Meyer, Xianshu Bai, Shan Bian, Anja Scheller, Frank Kirchhoff, Wenhui Huang

## Abstract

Molecular pathways mediating systemic inflammation entering the brain parenchyma to induce sepsis-associated encephalopathy (SAE) remain elusive. Here, we report that in mice during the first 6 hours of peripheral lipopolysaccharide (LPS)-evoked systemic inflammation (6 hpi), the plasma level of adenosine quickly increased and enhanced the tone of central extracellular adenosine which then provoked neuroinflammation by triggering early astrocyte reactivity. Specific ablation of astrocytic A1 adenosine receptors (A1ARs) prevented this early reactivity and reduced the levels of inflammatory factors (e.g., CCL2, CCL5, and CXCL1) in astrocytes, thereby alleviating microglial activation, ameliorating blood-brain barrier disruption, neuronal dysfunction, and depression-like behaviour in the mice. Chemogenetic stimulation of Gi signaling in A1AR-deficent astrocytes at 2 and 4 hpi of LPS injection could restore neuroinflammation and depression-like behaviour, highlighting astrocytes rather than microglia as early drivers of neuroinflammation. Our results identify early astrocyte reactivity towards peripheral and central levels of adenosine as a novel pathway driving SAE.

## Introduction

Severe systemic inflammation caused by infections or injuries can eventually deteriorate to a life-threatening status called sepsis (van der Poll et al., 2017). Such systemic inflammation also induces impaired functions of the central nervous system (CNS), one of the first affected organs in sepsis, leading to so-called sepsis-associated encephalopathy (SAE) (Mazeraud et al., 2020). Neuroinflammation is a hallmark of SAE, contributing to blood-brain barrier (BBB) damage, altered neuronal activity, and sickness/depression-like behaviour (Dantzer et al., 2008; Manabe and Heneka, 2022). Immune signals mediating systemic inflammation were shown to evoke rapid inflammatory responses firstly in components of the BBB (i.e., endothelial cells and pericytes of the brain vasculature) 1-2 h after the systemic inflammation challenge, subsequently releasing inflammatory mediators such as prostaglandin E2 or the chemokine (C-C motif) ligand 2 (CCL2, also referred to as monocyte chemoattractant protein 1, MCP1) and further potentiating neuroinflammation as well as aberrant neuronal activity (Dantzer et al., 2008; Fritz et al., 2016; Duan et al., 2018; Kodali et al., 2021). However, astrocytes were usually considered as targets of reactive microglia after a systemic inflammation challenge rather than early drivers of neuroinflammation themselves, which is counterintuitive concerning their anatomic positions as gatekeepers of the BBB easily receiving signals from the circulation.

Previous studies focused on augmented proinflammatory cytokines (e.g., TNFα, IL-1α and β, and IL-6) of the blood as immune signals which induce inflammatory responses in the CNS (Dantzer et al., 2008). In addition to cytokines, different purine metabolites such as ATP and adenosine can be found at elevated blood levels after a systemic inflammation challenge (Martin et al., 2000; Ramakers et al., 2011b; Sumi et al., 2014; Wu et al., 2022). However, the roles of purinergic signaling, if any, in affecting initiation and progression of SAE are largely undetermined. In the present study, we examined whether and how extracellular adenosine, the end-product of ATP hydrolysis, could regulate neuroinflammation using a well-established mouse model of SAE in which sepsis is induced by peripheral (intraperitoneal, i.p.) injection of the endotoxin lipopolysaccharide (LPS) (Liddelow et al., 2017; Duan et al., 2018; Kang et al., 2018; Hasel et al., 2021a). We detected rapid increases of extracellular adenosine levels in the blood and the brain shortly after the LPS injection which were followed by neuroinflammatory response of astrocytes, microglia and neurons in the CNS. We further showed the first *in vivo* evidence that elevated plasma adenosine could pass the BBB to enhance the central adenosine levels. Direct i.p. injection of adenosine or its analogues (i.e., 5′-N-ethylcarboxamide adenosine, NECA; and N6-cyclopentyladenosine, CPA) could induce an upregulation of several proinflammatory factors in the brain, which could be prevented by specific ablation of adenosine A1 receptors (A1ARs) in astrocytes rather than in pericytes, oligodendrocyte precursor cells or microglia, suggesting adenosine as an inflammatory mediator between the body and the CNS via triggering astrocyte reactivity. Comprehensive analyses of mice with inducible and astrocyte-specific A1AR conditional knockout (cKO) mice in the LPS-induced sepsis model further revealed that adenosine triggers astrocyte reactivity via A1ARs which provokes the inflammatory response of microglia at the early phase of systemic inflammation, driving the following pathogenesis of SAE by disrupting the BBB integrity, generating neuronal dysfunction as well as depression-like behaviour of the septic mice.

Therefore, we unveil a novel pathway employing adenosine as signaling molecule that mediates systemic inflammation to induce neuroinflammation by triggering early astrocyte reactivity. Moreover, we provide the first evidence that early reactive astrocytes act as drivers of neuroinflammation rather than being the targets of reactive microglia in SAE.

## Results

### Systemic inflammation increases adenosine levels in the blood and brain

Peripheral administration of a high dose of LPS (such as 5 mg/kg) is a widely used method to induce septic systemic inflammation in mice (Duan et al., 2018; Kang et al., 2018; Hasel et al., 2021a). Sepsis is known to increase plasma adenosine levels in patients or volunteers injected with low doses of LPS(Martin et al., 2000; Ramakers et al., 2011b). In the mouse model, we also found elevated levels of plasma adenosine at 2 hours post LPS injection (2 hpi), which peaked at 6 hpi, and subsequently dropped to baseline level at 12 and 24 hpi (Figures 1A and 1B). Previous studies suggested that systemic inflammation could disrupt BBB integrity(Haruwaka et al., 2019; Manabe and Heneka, 2022). In addition, adenosine itself acts on endothelial A1 and A2a ARs resulting in the opening of the BBB (Carman et al., 2011). Therefore, we performed Evans blue (EB) extravasation experiments to assess the BBB integrity after LPS injection. Mice were injected i.p. with EB two hours prior to each analysis time point (Figure 1A). Intriguingly, our experiments detected BBB disruption already at 2 hpi, which, similar to adenosine changes in the blood, reached the peak at 6 hpi and decreased at 12 and 24 hpi (Figure 1C). Furthermore, it has been shown that a peripheral injection of LPS could activate various glial cells and pericytes as well as several cytokines in the brain within 24 hours (Duan et al., 2018; Kang et al., 2018; Kodali et al., 2021). In our experiments, we also observed enhanced expression of genes related to inflammation (e.g., *Ccl2*, *Ccl5*, *Cxcl1*, *Cxcl10*, *Tnf*, *Il6*, *Il1a* and *Il1b*), astrocyte (*Gfap* and *Lcn2*) or microglia activation (*Itgam*) in the mouse cortex. Peaks of expression of the inflammation-related proteins were reached between 2-6 hours post injection (Figures S1A-1C), concomitant with a plasma adenosine increase and a BBB disruption, as signs of systemic inflammation.

**Figure 1.**
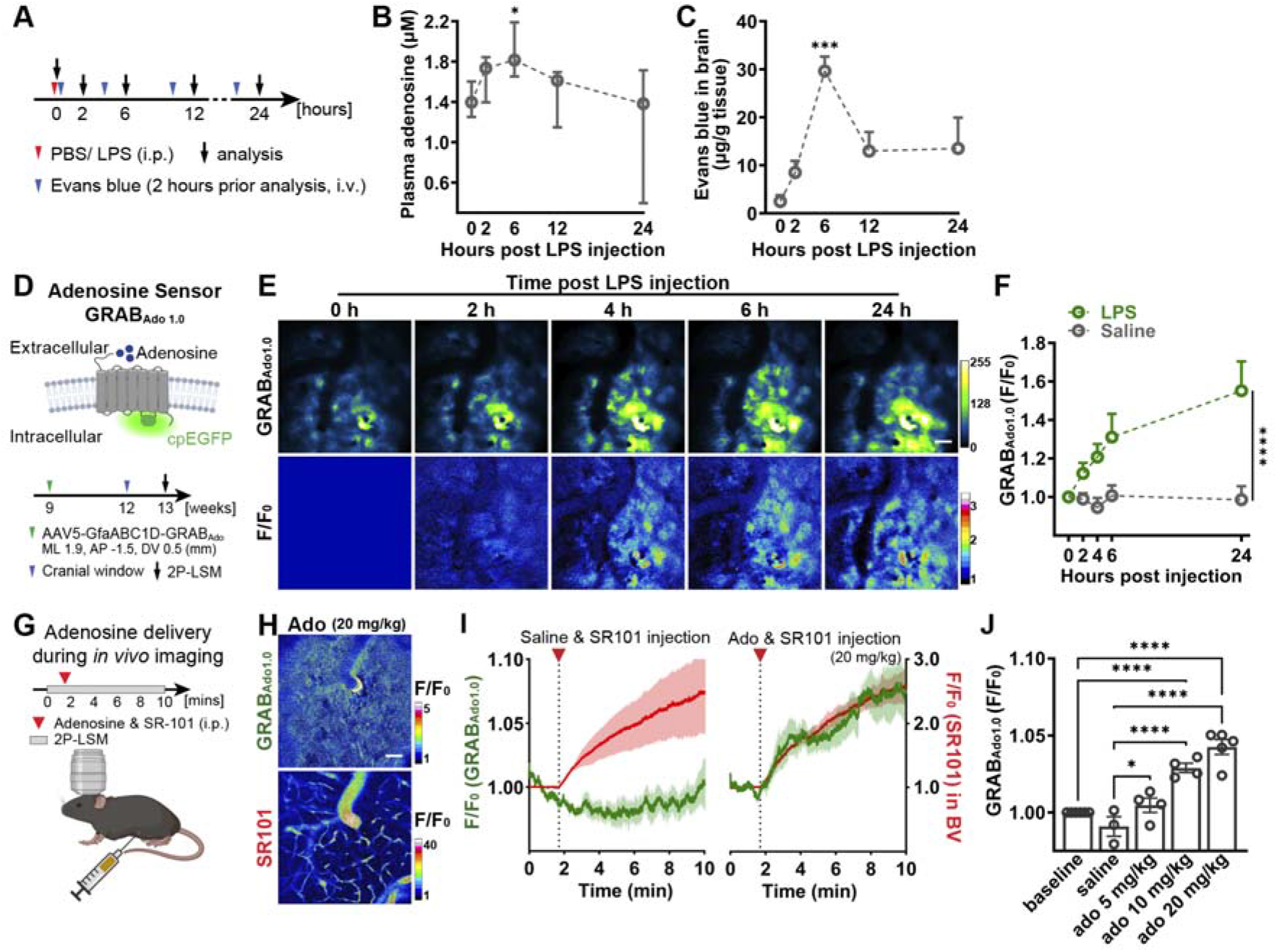
Peripheral LPS challenge increases adenosine levels in the brain and blood. **(A)** Schematic illustration of plasma adenosine level experiment **(B)** and brain Evans blue (EB) infiltration experiment **(C)**. **(B)** Plasma adenosine concentration rapidly increased post peripheral LPS injection (n = 5 mice for each time point). **(C)** EB extravasations increased in the brain after the peripheral LPS injection. EB was injected 2 h prior to the analysis time point (n = 3 mice for each time point). **(D)** Schematic illustration of the principle of the GRAB_Ado1.0_ sensors (up) and cortical extracellular adenosine level measurement post LPS injection by using GRAB_Ado1.0_ *in vivo* 2P-LSM live imaging (down). GfaABC1D-GRAB_Ado1.0_ plasmids were delivered to cortical astrocytes via AAV5 injection according to the coordinates indicated in the scheme. **(E)** Representative fluorescence images (up) and pseudocolor images (down) of GRAB_Ado1.0_ signals post peripheral LPS injection. Scale bar = 50□μm. **(F)** Comparison of relative fluorescence intensities (F.I.) of GRAB_Ado1.0_ acquired in the LPS/ Saline injection model. The recording of 0 h after LPS/Saline injection was used as F_0_. (n = 3 mice per group) **(G)** Schematic illustration of injection of adenosine supplemented with SR101 (i.p.) during the *in vivo* imaging of GRAB_Ado1.0_. **(H)** Representative pseudocolor images of GRAB_Ado1.0_ signals and SR101 signals after a peripheral adenosine and SR101 injection. Scale bar = 50□μm. **(I)** Increase of GRAB_Ado1.0_ signal (green) after the injection of adenosine (20 mg/kg, i.p.) was concomitant with the increase of SR101 signal (red) in blood vessels (BV), while GRAB_ado1.0_ signal is not altered after saline injection. **(J)** Relative fluorescence intensities (F.I.) of GRAB_Ado1.0_ upon applications of various dosages of adenosine. The baseline was used as F_0_. (baseline n = 6 mice, saline n = 3 mice, ado 5 mg/kg n = 4 mice, ado 10 mg/kg n = 4 mice, ado 20 mg/kg n = 5 mice) Summary data are presented as the median ± IQR (**B**) and mean ± SEM (**C**, **F**, **J**). Statistical significance in (**B**) was assessed using Kruskal-Wallis test; statistical significance in (**C**, **J**) were assessed by one-way ANOVA; statistical significance in (**F**) was assessed by two-way ANOVA, *P□<□0.05, **P□<□0.01, **** P□<□0.001. See also Figure S1.

To investigate whether systemic inflammation could also evoke extracellular adenosine level in the brain parenchyma, we performed *in vivo* 2-photon laser-scanning microscopy (2P-LSM) of a novel genetically encoded G protein-coupled receptor (GPCR)-activation-based (GRAB) sensor for adenosine (GRAB_Ado1.0_) (Figure 1D) (Peng et al., 2020) expressed in cortical astrocytes. We detected increased fluorescence intensities (F.I.) of GRAB_Ado1.0_ already 2 hpi, reaching plateau levels at 6 hpi (Figures 1E and 1F), suggesting that LPS-induced systemic inflammation could increase the extracellular adenosine level in the brain. Next, we examined whether peripheral adenosine in the blood could directly induce elevation of extracellular adenosine level in the brain. We injected i.p. adenosine (20 mg/kg) supplemented with sulforhodamine 101 (SR101, red fluorescence) to GRAB_Ado1.0_-expressing mice during 2P-LSM live imaging (Figure 1G). We were able to detect increased GRAB_Ado1.0_ F.I. after the adenosine injection, which is concomitant with the increase of SR101 F.I. in the blood vessels, while injection of SR101 alone could not increase the GRAB_Ado1.0_ F.I. (Figures 1H and 1I). In addition, we observed a dose-dependent increase of GRAB_Ado1.0_ F.I. when we injected different doses of adenosine (5 - 20 mg/kg, Figure 1J), indicating that increased peripheral adenosine can directly enhance the extracellular adenosine level in the brain. Taken together, although the potential sources of the increased extracellular adenosine in the brain after LPS treatment remain unidentified, our current results strongly suggest that peripheral LPS injection induced a rapid increase of plasma adenosine which may contribute to the elevated extracellular adenosine level in the brain as well as a rapid neuroinflammatory response in the early phase of the systemic inflammation.

### Peripheral adenosine administration induces astrocyte reactivity and neuroinflammation via A1ARs

To evaluate if adenosine could evoke a neuroinflammatory response, we injected adenosine (i.p., 5 mg/kg, six times, separated by 1 hour) to ‘wild-type’ (wt) control (ctl) mice and analysed gene expression levels in the mouse cerebral cortex by qPCR at 6 h post the first injection (Figure 2A). We injected adenosine several times due to its short lifetime in the blood (∼ 1 h)(Chiu et al., 2014).

**Figure 2.**
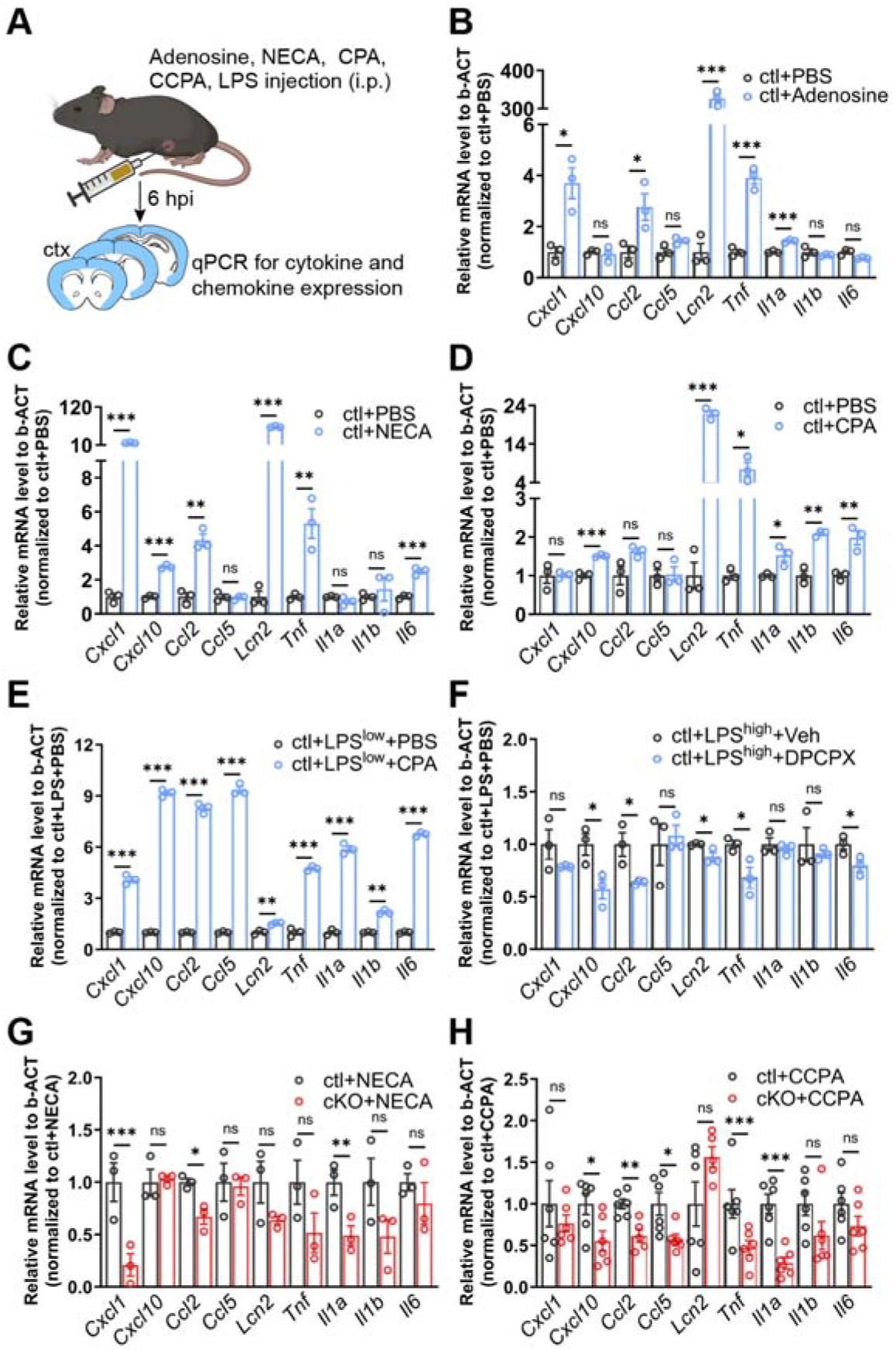
Peripheral adenosine administration evokes upregulation of inflammation-related genes in the brain. **(A)** Schematic illustration of adenosine, adenosine analogue (NECA), A1 adenosine receptor (A1AR) agonists (CPA, CCPA), and A1AR antagonist (DPCPX) administration experiments .**(B-D)** Expression of inflammation-related genes were enhanced in the mouse cortex six hours post adenosine **(B)**, NECA **(C)** and CPA **(D)** injections (n = 3 mice per group). **(E)** CPA further upregulated the inflammation-related genes in the cortex induced by a peripheral LPS^low^ (1 mg/kg, i.p.) injection. (n = 3 mice per group). **(F)** DPCPX administration reduced the inflammation-related genes in the cortex induced by a peripheral LPS^high^ (5mg/kg, i.p.) injection. (n = 3 mice per group). **(G, H)** Inflammation-related gene expressions were reduced in the cortex of cKO mice at 6 hours post NECA **(G)** and CCPA **(H)** injection compared to ctl mice (n = 3 mice per group in (**G**), n = 6 mice per group in (**H**)). Summary data are presented as the mean ± SEM. Statistical significance of each gene expression in (**B-H**) were assessed using unpaired Student’s t test, ns: not significant, *P < 0.05, **P < 0.01, *** P < 0.001. See also Figure S1 and S2.

Intriguingly, *Lcn2* (a marker for inflammatory reactive astrocytes)(Agnew-Svoboda et al., 2022) was upregulated (Figure 2B) upon adenosine administration. Moreover, several other proinflammatory cytokine/chemokine genes such as, *Ccl2, Cxcl1, Il1a* and *Tnf* also rapidly responded to the adenosine application with increased expression level (Figure 2B). Because the plasma adenosine level increased rapidly with the LPS injection and peaked at 6 hpi, we injected NECA (a non-selective adenosine analogue, half-life = ∼ 5 h)(Carman et al., 2011; Bynoe et al., 2015) to ctl mice and analysed cytokine expression at 6 hpi (Figures 2A). We observed that several cytokines such as *Cxcl1, Cxcl10, Ccl2, Tnf, Il6* as well as *Lcn2* were also upregulated upon NECA treatment (Figure 2C). In addition, we injected CPA (a selective A1AR agonist, half-life = ∼ 0.5 h) to ctl mice and also detected upregulated cytokines and other proteins related to inflammation/astrocyte activation at 6 hpi (Figures. 2A and 2D). Furthermore, when we combined the injection of CPA with a low-dose of LPS (1 mg/kg, LPS^low^), the expression levels of all tested cytokines or chemokines were strongly augmented compared to mice injected with LPS only (Figure 2E). On the other hand, in the cortex of high-dose (5 mg/kg, LPS^high^) LPS-challenged mice the administration of DPCPX (8-Cyclopentyl-1,3-dipropylxanthine, a selective A1AR antagonist) inhibited the activation of glial cells and neurons (Figures S1D and S1E) as well as reduced the expression of several inflammatory factors (e.g., *Ccl2, Tnf, Lcn2, Il6*, and *Cxcl10*) (Figure 2F). In conclusion, our results strongly suggest that A1AR signaling directly contributes to the induction of neuroinflammation.

Recent transcriptomic studies suggested that A1ARs are the most abundantly expressed ARs in astrocytes(Zhang et al., 2014; Ximerakis et al., 2019). To determine the contribution of astrocyte-specific A1ARs to adenosine-evoked neuroinflammation, we cross-bred GLAST-CreERT2 mice with temporal control of astrocyte-specific gene recombination(Motori et al., 2013) to floxed A1AR mice(Scammell et al., 2003), thereby generating inducible astrocytic A1AR cKO mice. In addition, we crossbred RiboTag mice (Sanz et al., 2009) enabling specific and direct purification of mRNAs (mRNA^RiboTag^) from Cre-expressing cells in cerebral cortices to avoid pitfalls inherent to cell-sorting procedures. In parallel, we used a genetically encoded Ca^2+^ indicator mouse line (Rosa26-LSL-GCaMP3, Rosa26-GCaMP3)(Paukert et al., 2014) to study Ca^2+^ activity in astrocytes after tamoxifen administration. We performed qPCR for *Adora1*^RiboTag^ expression as well as *ex vivo* slice recordings of Ca^2+^ response to CPA application to confirm a successful gene excision of *Adora1* in cKO mice (Figure S2). Subsequently, we injected NECA or CCPA (2-chloro-N6-cyclopentyladenosine, a more selective A1AR agonist) to cKO and ctl mice and found that the expression of several upregulated cytokines and chemokines (e.g., *Il1a, Ccl2, Cxcl1*) was significantly attenuated in cKO mice (Figures 2G and 2H), which was not observed in mice with specific ablation of A1ARs in pericytes, oligodendrocyte precursor cells and microglia (data not shown), providing additional evidence that adenosine could trigger neuroinflammatory response via astrocytic A1AR signaling.

### A1AR signaling augments inflammatory response of reactive astrocytes in the early phase of systemic inflammation

Upregulation of the immediate early gene *Fos* family is frequently used to indicate activation of glia and neurons(He et al., 2019; Nagai et al., 2019b). Therefore, we performed immunohistochemistry of c-Fos combined with the astrocyte marker Sox9 on mouse brain slices to investigate the time course of astrocyte activation after LPS (5 mg/kg, and for all the following experiments) injection. We found that the expression of c-Fos in astrocytes of ctl mice was unaltered at 2 hpi but was upregulated at 6 hpi and reduced to basal level at 24 hpi. However, the c-Fos expression in astrocytes in cortex and striatum was significantly inhibited by the deficiency of A1ARs at 6 hpi, indicating a critical early temporal window of the astrocytic response to a systemic inflammation (Figures 3A-3C). Next, we performed high-throughput RNA-seq to study the impact of A1AR-mediated astrocyte activation at the molecular level. We used the RiboTag approach to directly purify astrocytic translated mRNA (mRNA^RiboTag^) from the cortex for RNA-seq analysis (Figure S3A). We observed that astrocytes from ctl mice strongly responded to the peripheral LPS challenge in terms of dynamically up and down-regulating genes with time after LPS injection, in line with the previous reports(Hasel et al., 2021a). However, the overall gene expression in A1AR-deficient astrocytes were much less altered upon the LPS injection at 6 and 24 hpi compared to ctl mice (Figure 3D, Figure S3B, Figure S4). In total, comparing A1AR-deficient astrocytes to ctl astrocytes, we found 12 (4 up, 8 down) differentially expressed genes (DEGs) at 0 hpi (healthy controls), 185 (38 up, 147 down) DEGs at 6 hpi, and 2643 (1465 up, 1178 down) DEGs at 24 hpi (Figures S3C-S3E). Further Hierarchical Clustering analysis by K-means classified DEGs into 8 clusters (Figure 3D, Figures S4). At 6 hpi, 125 genes (cluster 1, 5) were upregulated in ctl mice including cytokines and chemokines (e.g., *Ccl2, Ccl3, Ccl5, Il1a, Il1b, Il6, Cxcl1, Cxcl10*, etc.), transcription factors that promote inflammation (e.g., *Nfkb1, Nfkb2, Stat3*, etc.), and previously identified marker genes of early responding reactive astrocytes such as interferon-responsive genes (e.g., *Ifit1, 2, 3*) and genes encoding proteins for antigen presentation (e.g., *H2-D1, H2-K1*)(Hasel et al., 2021a). However, in astrocytes of cKO mice the expression of those pro-inflammatory genes was inhibited. Of note, at 6 hpi several immediate early genes (e.g., *Fos, Jun, and Junb*; cluster 5) were also reactively expressed in ctl astrocytes as previously reported (Hasel et al., 2021a; Kodali et al., 2021) and were inhibited in cKO astrocytes, in agreement with the c-Fos immunohistochemistry results. Unlike genes in cluster 5 (e.g., *Fos, Jun, Junb*) that were rapidly down-regulated to baseline level at 24 hpi, on average genes in cluster 1 (e.g., *Ccl5, Ccl9, Cxcl10, Stat3*) were still expressed at high levels in ctl astrocytes but inhibited in A1AR-deficient astrocytes at 24 hpi. In clusters 2, 4, and 6, 918 genes were slightly dysregulated in both ctl and cKO astrocytes at 6 hpi but were either up- (cluster 6) or downregulated (cluster 2 and 4) in ctl astrocytes at 24 hpi. However, the expression of these genes in cKO astrocytes appeared to be slightly altered at 24 hpi. By Metascape pathway analysis we found that genes in these clusters are responsible for ‘DNA repair’ (e.g., *Fancm, Msh3, Xpa,* etc. in cluster 6), ‘neuronal system’ (e.g., *Gabrd, Mpdz,* etc. in cluster 2), and ‘cell projection assembly’ (e.g., *Fnbp11, Tppp, Nek1, etc.* in cluster 4). We also observed that many genes defined in previous studies as ‘A1 astrocyte’ (e.g., *H2-D1, H2-T23, Gbp2, Iigp1, Psm8,* ect.) and ‘A2 astrocyte’ (e.g., *Clcf1, Tgm1, Ptx3,* etc.) markers were decreased in cKO astrocytes at 24 hpi (Figure S3F). Therefore, we assume that this difference in cKO mice at 24 hpi could be regarded as the consequence of reduced astrocytic inflammatory response at 6 hpi which may ameliorate the global neuroinflammation afterwards (see below). For example, the GSEA-KEGG pathway analysis indicated that genes involved in astrocytic inflammatory response pathways (e.g., ‘NF-kappa B signaling pathway’, ‘JAK-STAT signaling signaling pathway’, ‘IL-17 signaling pathway, NOD-like receptor signaling pathway’, ‘TNF signaling pathway’, etc.) (Sofroniew, 2020; Han et al., 2021) were supressed in cKO mice at 6 hpi (Figures 3E and 3F). Similarly, GSEA-GO term (biological process, BP) analysis also demonstrated that genes annotated to inflammation-related GO terms (e.g., ‘interleukin-1 production’, ‘toll-like receptor signaling pathway’, ‘receptor signaling via JAK-STAT’, ‘I-kappaB/NF-kappaB signaling’, ‘inflammatory response, cytokine response’, ‘interferon-gamma production’, etc.) were suppressed in A1AR-deficent astrocytes at 6 hpi (Figures 3G, 3H). Furthermore, Metascape analysis for upstream transcriptional regulators revealed that the suppressed genes in A1AR-deficient astrocytes at 6 hpi were regulated by transcription factors (e.g., *Jun, Nfkb1, Cebpb, Stat3, Fos, etc.*) known to regulate downstream proinflammatory pathways of reactive astrocytes (Figure 3I)(Sofroniew, 2020; Han et al., 2021). Taken together, our results demonstrated that in the early phase after a peripheral LPS challenge adenosine triggers via A1ARs the inflammatory response of reactive astrocytes which may influence the subsequent progression of neuropathology.

**Figure 3.**
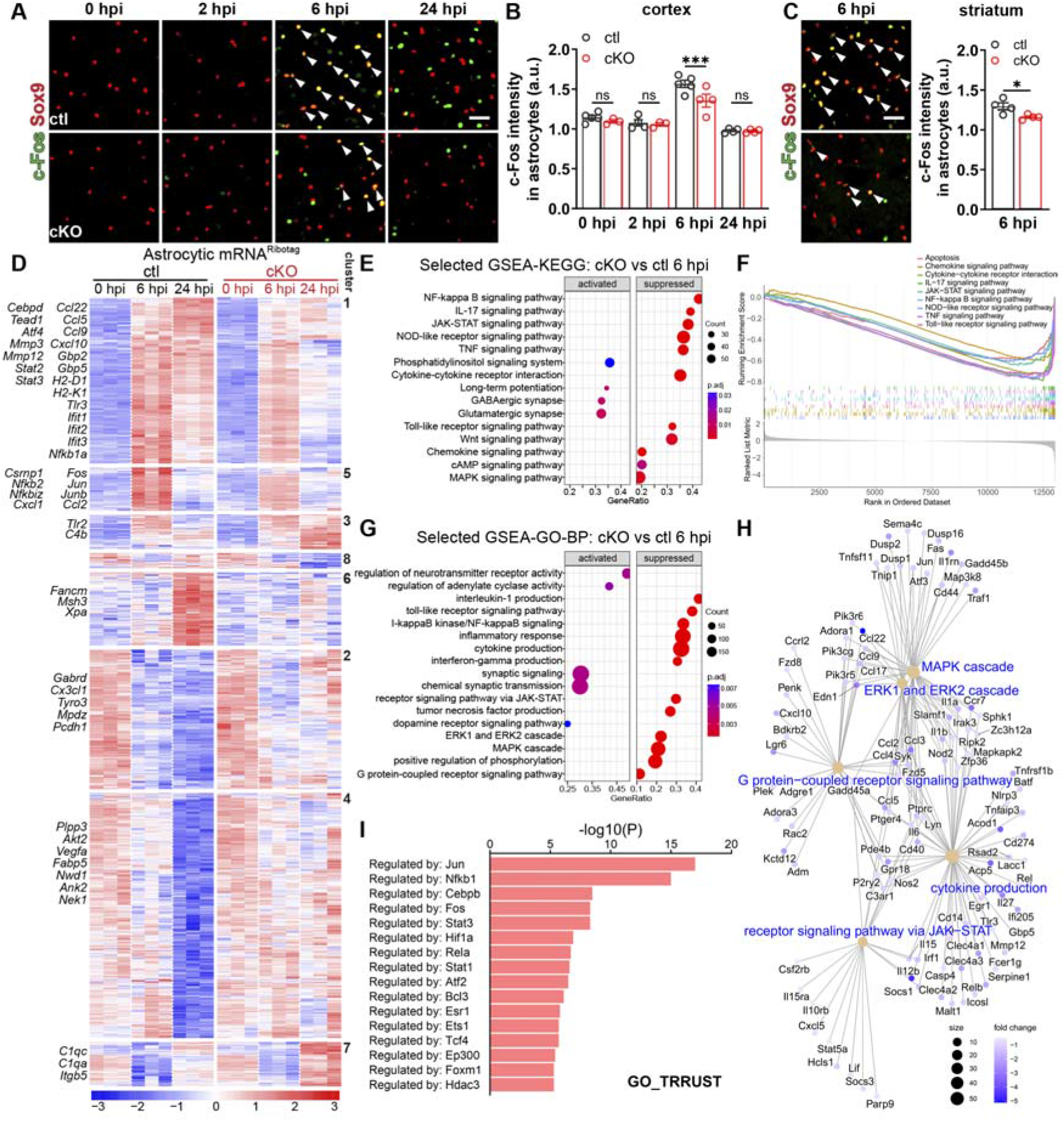
A1AR-deficient astrocytes are less reactive to the peripheral LPS challenge. **(A)** Representative images of c-Fos expression in cortical Sox9^+^ astrocytes (arrowheads) post LPS injection. Scale bar = 20□μm. **(B)** Cortical astrocytic c-Fos immunofluorescence intensity (arbitrary unit, a.u.) was enhanced at 6 hpi in ctl which was inhibitedin cKO mice(n = 5 mice in ctl at 0 hpi, n = 3 mice in cKO at 0 hpi, n = 4 mice in ctl at 2 hpi, n = 3 mice in cKO at 2 hpi, n = 4 mice in ctl at 6 hpi, n = 5 mice in cKO at 6 hpi, n = 6 mice in ctl at 24 hpi, n = 6 mice in cKO at 24 hpi). **(C)** Astrocytic c-Fos immunofluorescence intensity was enhanced at 6 hpi in the striatum of ctl which was reduced in cKO mice (n = 4 per group). Arrowheads indicated c-Fos^+^Sox9^+^ astrocytes. **(D)** Heatmap of altered gene expression (Padj < 0.05) from astrocytic mRNA^RiboTag^ of male cKO and ctl mice in any of the three time points (n = 3 mice per group). Clustering was done with 8 K-means. **(E)** Selected GSEA-KEGG pathway analysis of the astrocytic RNA-seq dataset between the cKO and ctl groups at 6 hpi. **(F)** Selected GSEA plot of the enriched KEGG pathways related to inflammation between the cKO and ctl groups at 6 hpi. **(G)** Selected GSEA-GO pathway analysis of the astrocytic RNA-seq dataset between the cKO and ctl groups at 6 hpi. **(H)** Category net plot of selected enriched GO pathways relative to G protein-coupled receptor signalling pathway. The color gradient indicates the fold changes between the cKO and ctl groups. **(I)** Prediction of transcription regulators following expression pattern of sub-clusters (cluster 1 and 5 in **(D)**) by Metascape analysis(Han et al., 2018; Zhou et al., 2019). Summary data of (**B**, **C**) are presented as the mean ± SEM. Statistical significance in (**B**) were assessed by two-way ANOVA; statistical significance in (**C**) were assessed by unpaired Student’s t test, ns: not significant, * P□<□0.05, *** P□<□0.001. See also Figure S3 and S4.

### Astrocyte reactivity in the early phase of systemic inflammation boosts the inflammatory response of microglia as well as global neuroinflammation

Microglia and astrocytes are key players in neuroinflammation. Previous studies suggested that astrocyte and microglia interact with each other to modulate neuroinflammation(Linnerbauer et al., 2020; Han et al., 2021). Particularly, in the peripheral LPS challenge model, astrocytes can become neurotoxic when triggered by cytokines released by activated microglia(Liddelow et al., 2017). However, these pioneer studies focused on the late phase (after 24 hpi) of the LPS model. Our current results show an adenosine-mediated early response (2-6 hpi) of astrocytes to LPS challenge that regulate the expression of inflammation-related genes, many of which (e.g., *Ccl2, Ccl5, Cxcl1 and Cxcl10*) have been shown to trigger the formation of reactive microglia(Sofroniew, 2020). Therefore, we hypothesized that astrocytes could modulate microglia activation in the initial phase of systemic inflammation rather than just be the effector of reactive microglia. It is known that reactive microglia produce many inflammatory cytokines via the NF-κB signaling pathway (Borst et al., 2021). Therefore, we first examined the reactivity of microglia in the early phase of the LPS model and detected nuclear translocation of p65 (a co-factor of NF-κB) as an indicator of reactive microglia (Figure 4A).

**Figure 4.**
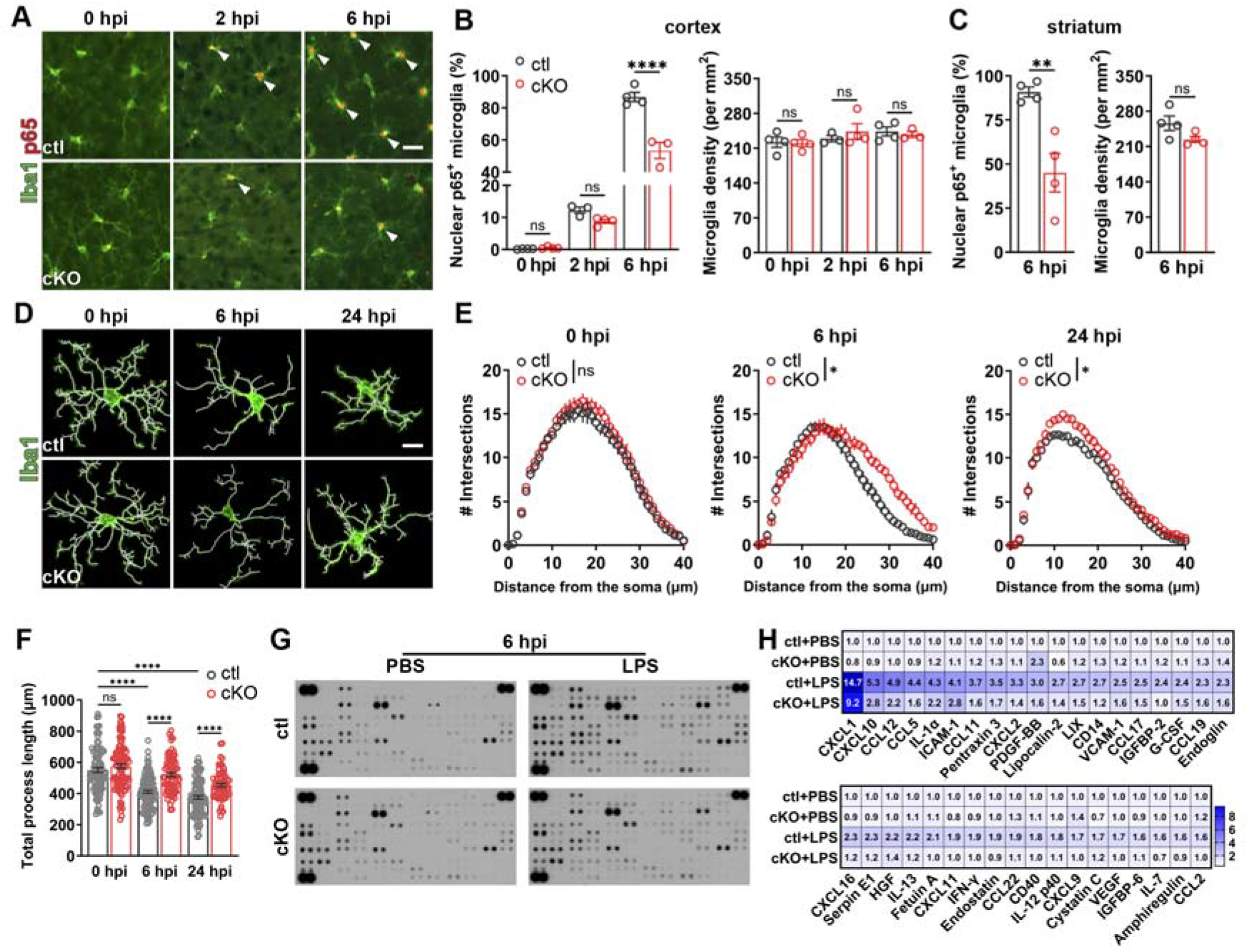
Astrocytic A1AR deficiency inhibits microglial activation and global neuroinflammation upon peripheral LPS challenge. **(A)** Representative images of p65 immunoreactivity in Iba1^+^ microglia post LPS injection. Arrowheads indicated Iba1^+^ microglia with nuclear P65. Scale bar = 20□μm **(B, C)** Proportions of nuclear p65^+^ microglia in the cortex (**B**) and striatum (**C**) of cKO mice were reduced post LPS injection compared to ctl mice, while the microglia densities were comparable in cKO and ctl mice (n = 4 mice in ctl/ cKO at 0 hpi, n = 3 mice in ctl at 2 hpi, n = 4 mice in cKO at 2 hpi, n = 4 mice in ctl at 6 hpi, n = 3 mice in cKO at 6 hpi in **B**; n = 4 mice per group in **C**). **(D)** Morphology of Iba1^+^ microglia post the LPS injection. 3D reconstruction was obtained using IMARIS. Scale bar = 10 μm. **(E)** Sholl analysis of Iba1^+^ microglia at 0 hpi, 6 hpi, 24 hpi (n = 3 mice per group). **(F)** Total process length of Iba1^+^ microglia in cKO and ctl mice post LPS injection obtained from the IMARIS-based morphological analysis (n = 3 mice per group). **(G, H)** Expression of 111 cytokines in the cortex of ctl and cKO mice was measured by a proteomic profiling assay at 6 h after PBS or LPS i.p. injection (samples from 3 mice were pooled for each group) (**G**). Cytokines with significant changes compared to ctl (PBS) group were shown in the heatmap (**H**). Color bar range is between 0.5 and 9.5, out range value was labelled with dark blue. Summary data are presented as the mean ± SEM. Statistical significance in (**B**, **E**, **F**) were assessed using a two-way ANOVA. Statistical significance in (**C**) is assessed using unpaired Student’s t test; ns: not significant, *P < 0.05, **P < 0.01, ****P < 0.0001.

We found that peripheral LPS challenge increased the proportion of nuclear p65^+^ microglia in the cortex of ctl mice from ∼ 13% at 2 hpi to ∼ 90% at 6 hpi, whereas such proportion in cKO mice was reduced to ∼ 9% and ∼ 50% at 2 and 6 hpi, respectively (Figures 4A and 4B). However, at 24 hpi virtually no nuclear p65 expression was detected in ctl or cKO mice (data not shown). Similarly, in the striatum the nuclear p65^+^ microglia were also reduced in the cKO mice at 6 hpi (Figure 4C). Of note, the densities of microglia in cKO and ctl mice were comparable (Figures 4B and 4C). Therefore, our results revealed that a systemic LPS challenge initiates astrocyte activation in the early phase via A1AR signaling to augment microglial activation. To further confirm this observation, we used IMARIS software to analyse microglial morphologies in cKO and ctl mice (Figure 4D). In the healthy control group (0 hpi), microglia of cKO and ctl mice displayed comparable morphologies. However, 6 or 24 hours after LPS injection microglia in cKO mice showed more intersections in the Sholl-analysis, more total process length, and a larger occupied area (Figures 4D-4F, Figure S6A), indicating reduced microglial activation in the absence of astrocytic A1AR signaling.

We next examined the impact of astrocytic A1AR signaling on the global inflammatory response of the brain following a systemic inflammation challenge. We measured 111 different cytokines in the cortex of ctl and cKO mice using a proteomic profiling assay and observed that most of the inflammation-related proteins in the cortex of ctl mice were upregulated after LPS injection at 6 and 24 hpi. However, in cKO mice the levels of several astrocyte specific/related proinflammatory proteins (e.g., CXCL1, CXCL10, ICAM-1, lipocalin-2 (LCN-2), MMP3)(Sofroniew, 2020) and many other cytokines or chemokines of multiple origins including microglia and astrocytes (e.g., CCL2, CCL5, IL-1a, CXCL2, etc.)(Borst et al., 2021) were largely reduced after LPS injection (Figures 4G and 4H; Figures S5C and S5D). In addition, the differentially expressed protein, LCN2 (Figure S6B), was chosen for Western blot to further confirm the results from the cytokine array. A similarly reduced inflammatory response pattern was detected in the striatum of cKO mice using a small-size (40 cytokines) cytokine array immunodot-blot assay (Figures S6E and S6F). Taken together, astrocytes were activated via A1ARs in the early phase of systemic inflammation to enhance microglial activation, exacerbating the global neuroinflammation afterwards.

### Astrocytic A1AR signaling contributes to impaired BBB function in systemic inflammation

Next, we investigated BBB function-related parameters in cKO and ctl mice following a peripheral LPS challenge. Systemic inflammation induces the CCL5-CCR5 axis-dependent migration of microglia to blood vessels, which further impairs the BBB(Haruwaka et al., 2019). We found that after LPS injection the perivascular microglia in ctl mice increased proportionally from ∼ 35% at 0 hpi to ∼ 50% at 6 and 24 hpi, while the proportion of perivascular microglia in cKO mice was only increased from ∼ 35% to ∼ 43% at 6 and ∼ 45% at 24 hpi, indicating a significant reduction (∼ 50%) of newly recruited perivascular microglia in cKO mice after peripheral LPS injection (Figures S5A-5C), in line with the reduction of CCL5 expression in the cKO mice. To assess the BBB integrity, we injected EB i.p. to mice immediately after the LPS injection and allowed EB to circulate for 24 hours. Afterwards, we measured EB extravasation in brain tissue. We detected less EB in the brain of cKO mice than in ctl (∼ 10 vs. 22 µg/g tissue, Figure S5D), indicating less disruption of the BBB in cKO mice. Systemic inflammation promotes infiltration of neutrophils into the brain parenchyma, attributable to the disrupted BBB as well as an upregulated expression of chemokines (e.g., CXCL1)(Huang et al., 2023) and extracellular matrix proteins (e.g., ICAM-1)(Rummel et al., 2010). Therefore, we performed Ly6B immunostaining to detect neutrophils. We found that the entry of Ly6B^+^ cells into the brain parenchyma was largely reduced 24 hours after the peripheral LPS injection in cKO mice compared to ctl (Figures S5E and S5F), in line with our current finding of reduced CXCL1 and ICAM-1 expression in cKO mice. Taken together, astrocytic A1AR deficiency ameliorated the systemic inflammation-induced impairment of the BBB.

### A1AR-mediated astrocyte activation promotes aberrant neuronal functions and depression-like behaviour in systemic inflammation

Systemic inflammation is also known to alter neuronal hyperactivity. For example, two hours after peripheral administration of LPS, pericytes are triggered to release CCL2, which subsequently determines neuronal hyperactivity(Duan et al., 2018). Therefore, we performed c-Fos and NeuN (a neuron marker) double immunostaining and studied neuronal activation after a peripheral LPS challenge. In contrast to astrocytes, we observed that c-Fos expression in cortical neurons in ctl mice gradually increased from 6 hpi to 24 hpi. Although c-Fos expression was still elevated in neurons from cKO mice compared with basal levels (0 hpi), it was lower than in ctl mice at 6 and 24 hpi (Figures 5A and 5B). Similarly, we detected lower c-Fos levels in striatal NeuN^+^ neurons of cKO mice at 24 hpi (Figure 5C). Thereby, these results further confirm that astrocytic A1ARs contribute to neuronal activation upon the challenge of systemic inflammation.

**Figure 5.**
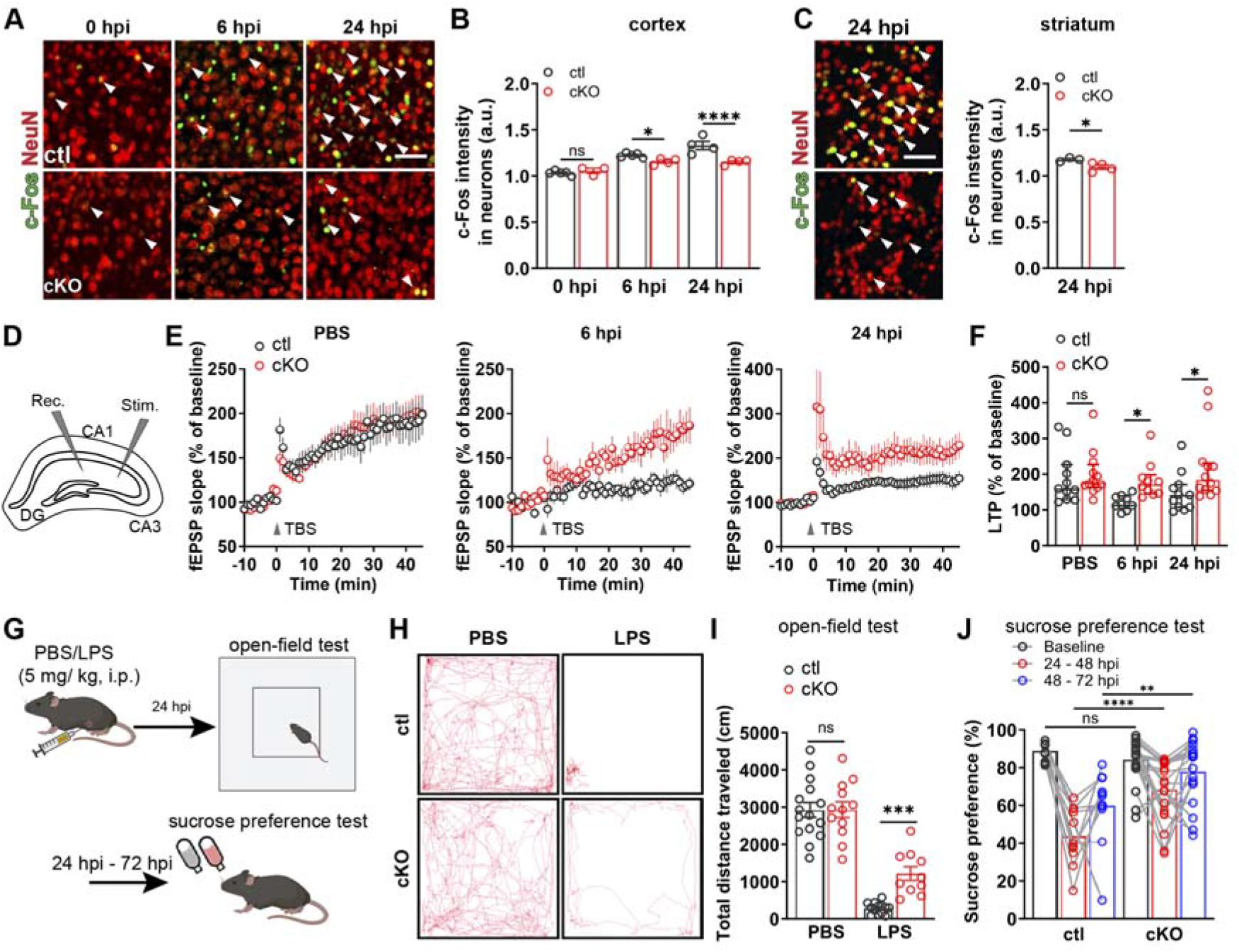
Astrocytic A1AR deficiency prevents neuronal dysfunction and ameliorates depression-like behaviour of the mice after LPS treatment. **(A)** Representative images of cFos immunoreactivity in cortical NeuN^+^ neurons after LPS injection. Scale bar = 20□μm **(B, C)** c-Fos immunofluorescence intensities (a.u.) in NeuN^+^ neurons in cortex (**B**) and striatum (**C**) of cKO mice were reduced compared to ctl mice at 6 and 24 hpi (n = 4 mice in ctl/ cKO at 0 hpi, n = 5 mice in ctl at 6 hpi, n = 3 mice in cKO at 6 hpi, n = 6 mice in ctl at 24 hpi, n = 4 mice in cKO at 24 hpi in **B**; n = 4 mice per group in **C**). **(D)** Graphical description of LTP measurement protocol. **(E)** Scatter plots showing LTP induced by stimulation of Schaffer collateral (SC) — *cornu ammonis* (CA) 1 synapses with TBS in acute hippocampal slices from ctl and cKO at 0 hpi, 6 hpi, 24 hpi. Averaged fEPSP are plotted versus time (n = 11 slices in ctl at 0 hpi, n = 12 slices in cKO at 0 hpi, n = 9 slices in ctl at 6 hpi, n = 10 slices in cKO at 6 hpi, n = 11 slices in ctl at 24 hpi, n = 12 slices in cKO at 24 hpi). **(F)** LTP evoked in hippocampi of cKO and ctl mice. **(G)** Schematic illustration of open-field test and sucrose preference test post PBS/LPS injection. **(H)** Representative trajectory analysis of ctl and cKO mice in 10 min in the open-field test at 24 hpi. **(I)** cKO mice displayed protected locomotion compared to ctl mice at 24 hpi (n = 15 mice in ctl PBS group, n = 12 mice in cKO PBS group, n = 12 mice in ctl LPS group, n = 10 mice in cKO LPS group). **(F) (J)** cKO mice displayed less LPS-induced decreased in sucrose preference than ctl after LPS injection (n = 11 mice in ctl group, n = 20 mice in cKO group). Summary data are presented as the mean ± SEM (**B, C, I, J**) and median ± IQR (**F**). Statistical significance in (**B, E, F, I, J**) were assessed using a two-way ANOVA; statistical significance in (**C**) was assessed using a unpaired Student’s t test. ns: not significant, *P < 0.05, **P < 0.01, ***P < 0.001, ****P < 0.0001.

Neuroinflammation induced by systemic endotoxin has been shown to affect neuronal network plasticity in terms of impaired long-term potentiation (LTP)(Shemer et al., 2020; Izumi et al., 2021; Wu et al., 2021). To study the impact of astrocyte-specific A1AR deficiency on neuronal plasticity following a peripheral LPS challenge, we performed extracellular electrophysiological recordings on acute slices prepared from dorsal hippocampi of cKO and ctl mice. Neurons from both groups displayed comparable LTP after PBS injection. However, LTP was impaired after LPS injection in ctl mice at 6 and 24 hpi, whereas it was preserved in cKO mice (Figures 5D-5F). These results indicate that adenosine triggers an astrocytic inflammatory response as a detrimental regulator of neuronal functions after systemic LPS challenge.

Neuroinflammation induced by peripheral LPS administration leads to sickness behaviour in mice (e.g., hunched posture, reduced food and water intake, etc.) that peaks between 2-6 hpi and gradually wanes. Subsequently, the depression-like behaviour (e.g., reduced locomotion in the open-field test, loss of preference to drink sweetened water, etc.) gradually develops to reach the peak 24 hours later(Dantzer et al., 2008; Kang et al., 2018). Therefore, we performed the open-field test to assess the general exploratory locomotion of mice as a readout of depression-like behaviour at 24 hpi. We observed ctl and cKO mice injected with PBS performed similarly in the open-field test. As expected, after LPS injection all ctl and cKO mice drastically reduced locomotion, whereas cKO mice moved longer distances than ctl mice (Figures 5G-5I), suggesting that the deficiency of astrocytic A1ARs could alleviate depression-like behaviour induced by systemic inflammation. In addition, we performed the sucrose preference test and assessed anhedonia, another hallmark of systemic inflammation-induced depression like behaviour (Figure 5G). We observed that peripheral LPS challenge could largely abolish the sucrose preference of ctl mice from 24-48 hpi (∼ 45%) which partially recovered to ∼ 60% from 48-72 hpi. However, the sucrose preference of cKO mice was only partially impaired to ∼ 65% from 24-48 hpi, and almost fully recovered to the healthy level from 48-72 hpi (∼ 80%), again indicating activation of astrocytic A1ARs contributes to depression-like behaviour induced by systemic inflammation (Figure 5J).

### Enhancing Gi signaling in A1AR-deficient astrocytes restored neuroinflammation upon peripheral LPS challenge

A1ARs are G_i/o_ protein-coupled receptors. Therefore, a weakened Gi signaling can be assumed in astrocytes of cKO mice. The Gi signaling could be artificially rescued using Designer Receptors Exclusively Activated by Designer Drugs (*DREADD*)-based chemogenetic tool hM4Di upon administration of its specific agonist CNO (clozapine N-oxide). Since previous studies highlighted that reactive microglia in the mouse striatum is a major contributor to depression-like behaviours after peripheral LPS challenge (Klawonn et al., 2021), we stereotactically injected AAV-GFAP-hM4Di-mCherry or AAV-GFAP-tdTomato into the striatum of cKO and ctl mice to specifically express hM4Di (indicated by mCherry) or tdTomato (tdT, as control) in astrocytes. To study whether A1AR-mediated astrocyte activation at the early phase of systemic inflammation is crucial to provoke the subsequent neuroinflammation, we injected CNO (half-life= ∼ 0.5 h) to AAV-delivered mice to evoke Gi signaling at 2 and 4 h post peripheral LPS challenge (Figures 6A-6C). At 6 hpi, we observed that in the cKO mice c-Fos expression was more enhanced in hM4Di-expressing rather than in tdT-expressing astrocytes, indicating that Gi signaling could further activate A1AR-deficient astrocytes post LPS injection (Figures 6D and 6E). Concomitantly, we found increased nuclear p65^+^ microglia in the regions with hM4Di-expressing astrocytes compared to tdT-expressing astrocytes in cKO mice, suggesting that enhancing astrocytic Gi signaling could promote reactive microglial inflammatory response at the early stage of systemic inflammation (Figures 6F and 6G). To assess the long-term effect of activation of Gi signaling in A1AR-deficient astrocytes, we measured the expression of proinflammatory factors in AAV-infected brain regions by cytokine array 24 h post LPS injection.

**Figure 6.**
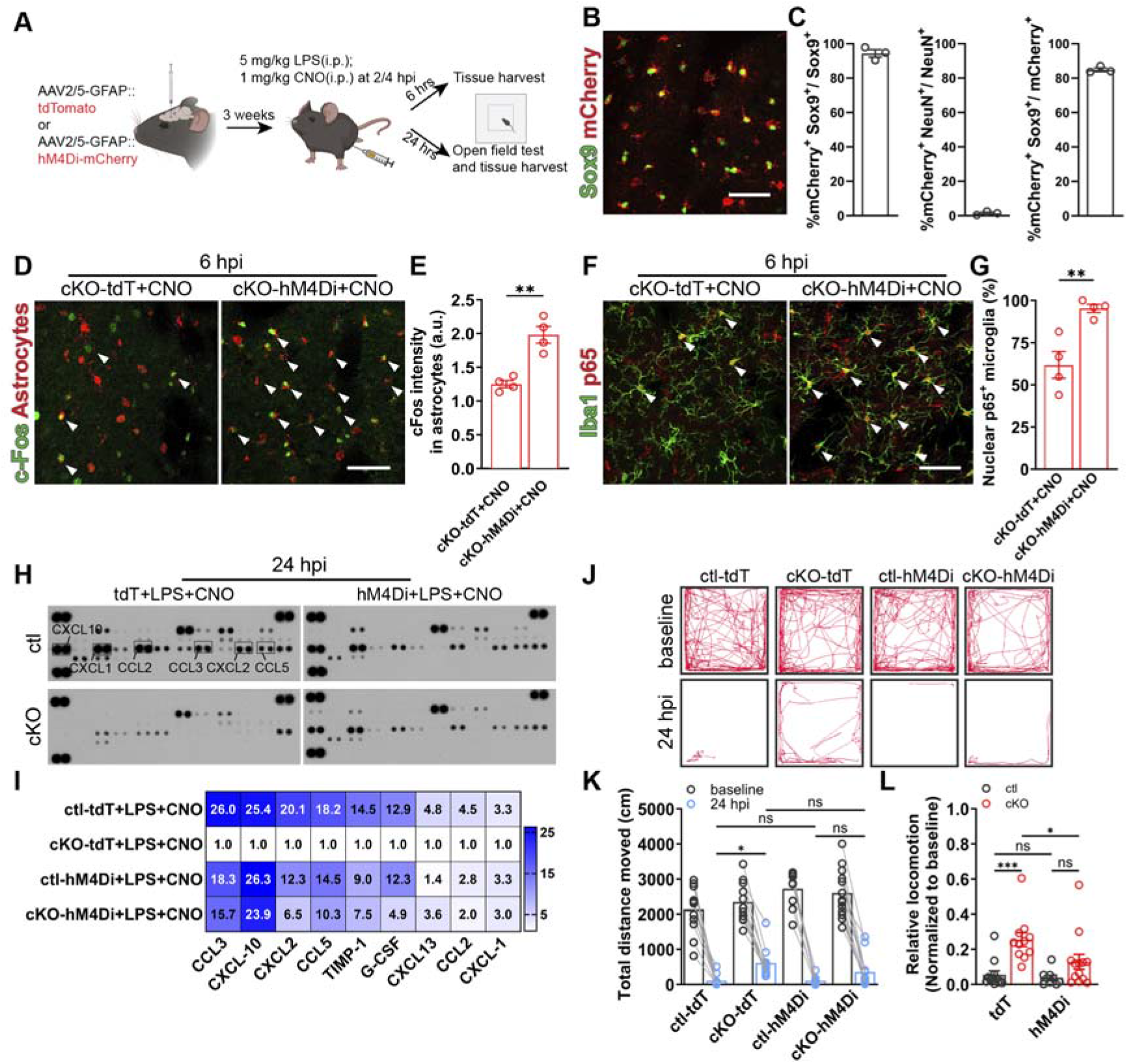
Activation of Gi signaling in A1AR deficit astrocytes restores the inflammatory response to peripheral LPS challenge. **(A)** Schematic illustration of activation of Gi signaling in A1AR deficit astrocytes experiment employing DREADD hM4Di and experimental plan. **(B)** Representative image of hM4Di expression indicated by mCherry in the Sox9^+^ astrocytes. Scale bar = 50□μm. **(C)** hM4Di was expressed in 94% of striatal astrocytes, with > 84% specificity (n = 3 mice). **(D)** Representative images of immunolabeled reactive A1AR-deficient astrocytes by c-Fos immunostaining after LPS and CNO injection. Scale bar = 50□μm. **(E)** Activation of hM4Di increased c-Fos expression in hM4Di-mCherry^+^ A1AR-deficient astrocytes after LPS injection (n = 4 mice per group). **(F)** Representative images of immunoreactivity of p65 in Iba1^+^ microglia in cKO mice with astrocytic tdT or hM4Di expression after LPS and CNO injection. Scale bars = 50□μm. Arrowheads indicates Iba1^+^ microglia with nuclear p65 expression. **(G)** Nuclear p65^+^ microglia was increased in hM4Di-expressing cKO mice after LPS and CNO injection compared to ctl mice (n = 4 mice per group). **(H)** The expression of 40 cytokines in the striatum of AAV-infected ctl and cKO mice was measured by a proteomic profiling assay after LPS and CNO injection. **(I)** Enhancing Gi signaling in cKO mice increased cytokine expression after LPS and CNO injection (samples from 3 mice were pooled for each group). **(J)** Representative trajectory analysis digitally tracked movement of ctl and cKO mice injected with LPS and CNO in the open-field test at 24 hpi. **(K, L)** Enhancing Gi signaling in A1AR-deficit astrocyte reduced locomotion in the open-field test (n = 12 mice in ctl-tdT+CNO group, n = 11 mice in cKO-tdT+CNO group, n = 9 mice in ctl-hM4Di+CNO group, n = 13 mice in cKO-hM4Di+CNO group). Summary data are presented as the mean ± SEM. Statistical significance in (**E**, **G**) were assessed using unpaired Student’s t test; statistical significance in (**K, L**) was assessed using a two-way ANOVA, ns: not significant, *P < 0.05, **P < 0.01, ***P < 0.001.

We observed that the expression of the factors (e.g., CCL2, CCL5, CCL3, CXCL-1, CXCL2, CXCL10, G-CSF, etc.) inhibited in the tdT-expressing cKO mice (compared to tdT-expressing ctl mice) could be enhanced to certain extents in hM4Di-expressing cKO mice (Figures 6H and 6I). In addition, a similar effect could be detected in the cortex of cKO mice with astrocytic hM4Di activation (Figures S6G and S6H), suggesting a consistent effect of Gi signaling in astrocytes from different brain regions to promote neuroinflammation. Also, we performed an open-field test on the same mice and found that the alleviated depression-like behaviour in cKO mice upon LPS challenge could be exacerbated to the level of ctl mice upon astrocytic Gi signaling activation (Figures 6J-6L). In summary, our results strongly suggest that early astrocyte activation via A1AR signaling in systemic inflammation plays a pivotal role in promoting neuroinflammation to drive pathogenesis of SAE.

## Discussion

Adenosine has been widely recognized as a guardian molecule that executes protective function in and outside of the CNS. For example, in the periphery adenosine reduced inflammation by triggering A2a, A2b, and A3 ARs of immune cells (e.g., macrophages, lymphocytes, neutrophils, etc.) as demonstrated, among others, in autoimmune diseases (e.g., rheumatoid arthritis), inflammatory bone loss, intestinal inflammation(Borea et al., 2016). In the CNS, adenosine is known to trigger pre- and post-synaptic A1ARs to reduce neuronal activity and, thereby, to alleviate epilepsy and pain(Borea et al., 2016). However, recent studies also suggest that the overproduction of adenosine could enhance neuroinflammation, for example, via microglial A2a ARs in Parkinson disease (PD)(Borea et al., 2017; Meng et al., 2019). In the current study, we revealed that peripheral administration of adenosine or its analogues can directly trigger inflammatory responses in the brain mediated at least partially by astrocytic A1ARs. Using peripheral LPS challenge, we provide further evidence that systemic inflammation-induced adenosine rise in the brain plays a pivotal role via astrocytic A1AR signaling in exacerbating neuroinflammation to drive the pathogenesis of the SAE. Therefore, like its precursor ATP, adenosine can also behave like a DAMP (damage-associated molecular patterns) molecule for neuroinflammation, possibly in a context-dependent manner.

Adenosine is mainly formed from the sequential ATP metabolism by means of a series of hydrolases, among which the ecto-5’-nucleotidase Nt5e (also known as CD73) performs the last step converting AMP to adenosine(Boison, 2008; Welsh and Kucenas, 2018). Under cellular stress conditions, such as inflammation or hypoxia, extracellular ATP/adenosine can drastically increase(Benarroch, 2008; Chiu and Freund, 2014). Recent studies using a novel genetically modified ATP sensor (GRAB_ATP1.0_) provided direct evidence that extracellular ATP levels in the mouse brain were increased after peripheral LPS challenge(Wu et al., 2022). Therefore, it is conceivable that increased extracellular ATP in the brain may contribute to adenosine elevation as well. Indeed, here we detected elevated extracellular adenosine in the brain parenchyma upon LPS injection employing *in vivo* 2P-LSM imaging of GRAB_Ado1.0_ and this effect lasted at least 24 h post the LPS challenge. In addition, adenosine can be transported by equilibrative nucleoside transporters (ENTs) or can bypass, in virtue of its small size, tight junctions (TJs) of endothelial cells(Latini and Pedata, 2001; Mikitsh and Chacko, 2014). Furthermore, adenosine can increase the permeability of the BBB through A1 and A2a ARs in endothelial cells, which facilitate the infiltration of peripheral immune cells (or even bacteria) expressing CD73 into the brain parenchyma(Mills et al., 2008; Carman et al., 2011; Bynoe et al., 2015; Zhao et al., 2020). To determine whether humoral adenosine could not only open the BBB but also further diffuse into the brain parenchyma, we injected adenosine i.p to mice during the 2P-LSM recording of GRAB_Ado1.0_ signals. We were able to detect increasing GRAB_Ado1.0_ signals in the brain along with the entry of the peripherally administered adenosine into cerebral blood flow indicated by SR101 in a dose-dependent manner. Although locally generated extracellular adenosine was reported to be quickly eliminated within seconds, previous studies have shown that i.p. injected adenosine could remain in the blood for at least one hour(Chiu et al., 2014; Bynoe et al., 2015). Therefore, our 2P-LSM results of adenosine administration strongly suggest that peripherally administered adenosine could either pass through the BBB or trigger endothelial cells to release ATP/adenosine (or both), thereby increasing the extracellular adenosine level in the brain parenchyma. We also demonstrated (as found in septic patients and volunteers receiving LPS injection(Martin et al., 2000; Ramakers et al., 2011a)) that the plasma level of adenosine was increased by the LPS challenge gradually peaking at 6 hpi, concomitant with the progression of astrocytic c-Fos expression, BBB breakdown and sickness behaviour of the mice. Moreover, considering that the BBB integrity was impaired by systemic inflammation, which could permit even more adenosine entry from circulation to the brain parenchyma, it is conceivable that systemic inflammation-increased humoral adenosine adds to locally generated adenosine to affect cells building the BBB (e.g., astrocytes and pericytes), participating in neuroinflammation.

In line with recent studies(Hasel et al., 2021a), our c-Fos immunohistochemistry as well as RNA^RiboTag^ sequencing results revealed that astrocytes reacted rapidly within hours to a systemic inflammation challenge and underwent dynamic transcriptomic changes. Although many molecules such as TNFα, IL-1α from reactive microglia, ATP, NO, etc. have been identified as triggers of astrocyte reactivity(Sofroniew, 2020), the exact molecules that initiate/boost such fast astrocyte reactivity to systemic inflammation are still undefined(Patani et al., 2023). Here, we demonstrated that the removal of A1ARs from astrocytes could largely prevent such fast astrocyte reactivity upon LPS injection. Moreover, adenosine and its analogues were able to elicit a neuroinflammatory response in healthy mice, that was strongly attenuated by lack of astrocytic A1AR. Our results suggest that the formation of reactive astrocytes in the early phase of systemic inflammation requires proinflammatory cytokines from the circulation or other CNS cells, but small molecules from the metabolism could play important roles as well. Here, adenosine is proposed as the first identified small molecule. It triggers the early astrocytic response to systemic inflammation via A1AR signaling.

Recent transcriptomic studies demonstrated astrocytes and microglia are both fast responders at the early phase of systemic inflammation(Shemer et al., 2020; Hasel et al., 2021b; Kodali et al., 2021). Although reactive astrocytes were usually considered as the long-term effector of reactive microglia especially in the systemic inflammation model(Patani et al., 2023), whether this is the case for the early astrocyte reactivity is unclear. Our astrocytic RNA^RiboTag^ sequencing results revealed that astrocytes at 6 hpi were triggered to release a series of inflammatory factors that are known to activate microglia via NF-κB activation, which were largely inhibited upon the ablation of A1AR signaling. Compensating the downstream Gi signaling by DREADD hM4Di strategy in the A1AR-deficient astrocytes before 6 hpi could largely restore the NF-κB activation of microglia and global neuroinflammation. Therefore, we provide evidence that astrocytes at the early phase of the systemic inflammation behave more like a booster of reactive microglia. Furthermore, the immune signal from the inflamed circulation was shown to be relayed by cells building the BBB (i.e., endothelial cells and pericytes) to the brain parenchyma (Fritz et al., 2016; Duan et al., 2018; Kodali et al., 2021). As another major component of the BBB, each astrocyte extends its end-feet to wrap at least one blood vessel(Hösli et al., 2022). Our results suggest that, in addition to endothelial cells/pericytes, immune signals mediated by adenosine from the circulation stimulate astrocytes to subsequently enhance microglia activation at the initial phase of the systemic inflammation. Thereby, reactive microglia may in turn modulate astrocyte reactivity in terms of the temporally changed transcriptome towards the neurotoxic A1-type in the late phase of systemic inflammation (24 hpi). However, such adenosine-regulated reciprocal interactions between astrocytes and microglia needs further investigation in other neuroinflammatory diseases such as multiple sclerosis (MS). In addition, future studies need to determine the dynamics of the interactome between astrocytes and microglia (and other inflammation-related cells such oligodendrocyte precursor cells) to gain novel insights into the progression of neuroinflammation in sepsis.

In conclusion, we provide evidence that adenosine, a ubiquitous signaling molecule, behaves as an immune mediator of systemic inflammation to exacerbate neuroinflammation via astrocytic A1Ars, thereby driving the pathogenesis of SAE. Since A1ARs are the most abundant ARs expressed in the brain, understanding how A1Ars, which are expressed in other cell types (i.e., pericytes, microglia, oligodendrocyte precursor cells) as well, contribute to neuroinflammation will enhance our knowledge on adenosine signalling for inflammation-related CNS diseases and might be the basis for future therapeutical approaches.

## Acknowledgement

We thank Daniel Schauenburg and colleagues for excellent animal husbandry. We thank Dr. Yulong Li (Peking University) for providing the AAV-GRAB_Ado1.0_ plasmids; Dr. Amit Agarwal (University of Heidelberg) for providing Rosa26-GCaMP3 mice, Dr. André Zeug (Hannover Medical School) for providing the AAV2/5-GFAP-tdTomato virus. Schematic drawings were generated by BioRender. This work was supported by grants from the Deutsche Forschungsgemeinschaft (HU 2614/1-1 (Project No. 462650276) to W.H., Sino-German joint project KI 503/14-1 to W.H. and F.K., SPP 1757, SFB 894 to F.K.); the Fritz Thyssen Foundation (10.21.1.021MN) and the Medical faculty of the University of Saarland (HOMFORexzellent2016) to W.H.; the BMBF (EraNet-Neuron BrIE) and the European Commission (EC-H2020 FET ProAct Neurofibres) to F.K.. Q.L. was supported by a PhD student fellowship of the Chinese Scholarship Council.

## Author contributions

W.H. conceptualized the project and designed experiments. W.H. and F.K. supervised the study. Q.G. performed RiboTag immunoprecipitation, immunohistochemistry, immunoblot, qPCR, confocal imaging and analysis, RNA-seq analysis, and behavioral test. Q.G. and D.G. performed AAV injection. Q.G., D.G., and W.H. carried out *in vivo* 2P-LSM imaging. D.G. performed *ex vivo* slice imaging and all 2P-LSM imaging analysis. N.Z. performed LTP recordings. Q.G., T.G. and M.R.M performed plasma adenosine concentration measurement. Q.L. contributed to immunohistochemistry and behavioral tests. L-P.F., X.B. and S.B. contributed to sample collection and qPCR. A.S. performed AxioScan imaging. Q.G. analyzed data and generated figures. W.H. and Q.G. wrote the manuscript with input from all the other authors.

## Materials and Methods

### Animals

This study involved the following animals maintained on C57BL/6 background: A1AR^fl/fl^ mice(Scammell et al., 2003); Glast-CreERT2(CT2) mice(Mori et al., 2006); RiboTag mice (Rlp22HA) (Sanz et al., 2009); GCaMP3 reporter mice (Rosa 26-CAG-lsl-GCAMP3)(Paukert et al., 2014). To specifically delete A1AR in astrocytes, A1AR^fl/fl^ mice were crossed to Glast-CreERT2 mice to generate A1AR^fl/fl^xGlast^ct2/wt^ mice, which were used as astrocytic A1AR conditional knockout mice (cKO) upon tamoxifen administration. For Figure 1, Figures 2B-F, and Figure S1, C57BL/6, A1AR^wt/wt^xGlast^ct2/wt^ and A1AR^fl/fl^xGlast^wt/wt^ littermate mice were used as ‘wild-type’ (wt) control (ctl) mice without tamoxifen injection. For other experiments, A1AR^wt/wt^xGlast^ct2/wt^ and A1AR^fl/fl^xGlast^wt/wt^ littermate mice were used as control (ctl) mice, also receiving tamoxifen injection.

RiboTag mice (Rlp22HA) were introduced to immunoprecipitate ribosome-associated translated mRNA in astrocytes upon breeding with A1AR^fl/fl^xGlast-CreERT2 mice. To visualize Ca^2+^ activity in astrocyte, GCaMP3 reporter mice were introduced to A1AR^fl/fl^xGlast-CreERT2 mice.

Mice of both sexes with 11-13 weeks age were used for most experiments (details in supplementary Table S1). All animals were maintained at the animal facility of the CIPMM under temperature- (22°C ± 2°C) and humidity- (45–65%) controlled conditions with 12□h light/dark cycle with an ad libitum supply of food and water. Animal husbandry and procedures were performed at the animal facility of CIPMM, University of Saarland according to European and German guidelines for the welfare of experimental animals. Animal experiments were approved by the Saarland state’s ‘‘Landesamt für Gesundheit und Verbraucherschutz” in Saarbrücken/Germany (animal license number: 65/2013, 12/2014, 34/2016, 36/2016, 03/2021 and 08/2021).

### Tamoxifen (TAM), LPS treatment

As previously described(Liu et al., 2023), tamoxifen (CC99648, Carbolution) was dissolved in Miglyol (3274, Caesar & Loretz, Hilden) at a concentration of 10 mg/ml and injected intraperitoneally to all mice crossed with CreERT2-driver mouse lines (100 mg/kg per body weight) for five consecutive days at the age of 4 weeks.

For the endotoxin challenge, 13 weeks old mice were injected intraperitoneally (i.p.) with 5□mg/kg of LPS (L2880, E. coli O55:B5, Sigma) or endotoxin-free PBS between 1 p.m. to 3 p.m. (CET) and kept for a maximum of 72 h post injection(Hasel et al., 2021b).

### Plasma adenosine measurement

To measure plasma adenosine level, 0.4 ml blood was collected from the right ventricle of deeply anesthetized mice with a syringe containing 0.4 ml ice-cold stop solution (220 μM APCP, 100 μM NBMPR, 40 μM dipyridamole, 13.2 mM Na2EDTA, 118 mM NaCl, 5 mM KCl), to block adenosine formation and transport. After centrifugation (1000g, 4°C, 10 min), plasma was stored at −80°C until analysis.

The plasma adenosine concentration was quantified with an adenosine assay kit (Fluorometric, Abcam) according to the protocol. The fluorescence was measured at excitation/emission = 535/587 nm.

### Evaluation of blood brain barrier (BBB) permeability by Evans blue

EB (4% wt/vol in saline, 2 mL/kg) was administered through the tail vein 2 h before mice were euthanized. Afterwards, Mice were perfused through the left ventricle with ice-cold PBS to remove intravascular albumin-Evans blue. Then half of the brains were quickly removed, weighed, and homogenized in 1 mL of PBS. After the first homogenization, 1 mL of 50% trichloroacetic acid were added, followed by vortex mixing for 2 min. After centrifugation (1000g, 4°C, 30 min), the supernatant was collected, and fluorescence intensity was measured at 620 nm on a microplate fluorescence reader and compared against a standard curve (serial dilutions of the stock dye solution in the concentration range of 0, 1.25, 2.5, 5, 10, 20 μg/mL). The Evans blue content was calculated and expressed as per gram of brain tissue.

### Stereotactic adeno-associated viruses (AAV) injection

10-week-old mice were used for AAV injection as described before with some modifications (Nagai et al., 2019a). Briefly, mice were administered 0.1 mg/kg of buprenorphine (s.c.) subcutaneously and 0.2 mg/kg of dexamethasone (i.p.) before surgery. Subsequently, under continuous inhalational isoflurane anesthesia (5% for induction and 2% for maintenance with 66% O_2_ and 33% N_2_O), the mouse head was fitted and secured by blunt ear bars in a stereotactic apparatus (Robot stereotaxic, Neurostar) and the mouse eyes were covered by Bepanthen (Bayer, Leverkusen). After sterile cleaning and skin incision, viruses were injected at a rate of 0.1□μl□min^−1^ with a 10 µl Nanofil syringe (34 GA blunt needle, World Precision Instruments) into the striatum (1.1 mm anterior to bregma, ±1.2 mm from sagittal suture at a depth of 3 mm from the skull surface) or cortex (1.9 mm posterior to bregma, 1.5 mm from sagittal suture at a depth of 0.5 mm from the skull surface). The syringe was kept in place for 10 min after the injection was completed, to avoid liquid reflux. The skin was closed by simple interrupted sutures (non-absorbable, F.S.T.). Carprofen and dexamethasone were administered once per day for three consecutive days and received either buprenorphine (1 mg/kg) or tramadol hydrochloride in the drinking water (400 mg/L) after surgery. Mice were used for experiments 3 weeks post virus injection. Viruses used were: AAV2/5 GfaABC1D-GRAB_Ado_ virus (2.3 × 10^12^ genome copies/ml); AAV2/5 GFAP-hM4Di-mCherry virus (3.04 × 10^12^ genome copies/ml); AAV2/5 GFAP-tdTomato virus (1.1 × 10^13^ genome copies/ml), 0.8 μl for striatum, 0.5 μl for cortex, respectively.

### Adenosine, adenosine analogue NECA, A1AR agonist CPA/CCPA, A1AR antagonist DPCPX treatment

To pharmacologically activate adenosine signaling, adenosine (5 mg/kg per body weight) was injected intraperitoneally every hour till 6 hours post first injection; NECA (1 mg/kg) was injected intraperitoneally once at 0 hpi; CPA (1 mg/kg) was injected intraperitoneally once at 0 hpi; CCPA (0.3 mg/kg) was injected intraperitoneally once at 0 hpi; DPCPX (1 mg/kg) was injected intraperitoneally 4 hours after LPS (5□mg/kg) or Veh (1% DMSO in PBS) injection. To further amplify inflammatory response, CPA (1 mg/kg) was injected intraperitoneally 4 hours after LPS (1□mg/kg). Mice were analyzed at 6 hours post injection.

### Craniotomy and window implantation

The skull of the mice was exposed and underwent a craniotomy followed by removal of the dura (Cupido et al., 2014; Kislin et al., 2014) in correspondence of the primary somatosensory cortex (SSp-ctx; Ø 3-4 mm, centered 1.5-2 mm posterior to bregma, 1.5 mm from the sagittal medial axis). The glass coverslip (Ø 3 mm, No. 1.5, Glaswarenfabrik Karl Hecht GmbH) and a custom-made holder for head restraining were fixed with dental cement (RelyXTM Unicem 2 Clicker, 3M Deutschland GmbH). Animals received canonical post-surgical care; three-day habituation training and imaging was performed one week after cranial window implantation.

### *In vivo* imaging of extracellular adenosine level

*In vivo* imaging from GRAB_ado_ virus injected mice was performed using a custom-made two-photon laser-scanning microscope (2P-LSM) setup. Recordings started one week after cranial window implantation to enable animal recovery and habituation according to adapted protocols without water restriction (Cupido et al., 2014; Kislin et al., 2014). During image acquisition at different time points after an LPS challenge, awake mice were maintained head-fixed and placed into a smooth plastic tube to reduce vibration caused by the movement of the mice. In the drug administration experiments, the head-fixed mice received 1.5% isoflurane anesthesia delivered by means of a custom-made nose mask. Solutions of adenosine (0, 5, 10 or 20 mg/kg) and SR101 (20 mg/kg) was prepared in isotonic saline solution and administered after 200 frames of baseline recording (∼ 88 s). Squared FOVs (512 x 512 μm) were chosen at a 60-80 μm depth from the surface and displaying a uniform virus expression with 910-nm laser. The laser power under the objective was adjusted from 5 to 30 mW, depending on the focal plane depth. Single focal plane images were acquired with a 2.28 Hz frame rate (1.4 µs pixel dwell time) and a resolution of 512 x 512 pixel.

### Reverse transcription-polymerase chain reaction (RT-PCR)

Deeply anesthetized mice were perfused with ice-cold PBS. Cortex and striatum were dissected in ice cold PBS using an adult brain reference atlas. Total RNA of cortex and striatum were isolated with the RNeasy mini kit (74106, Qiagen) according to the instructions and underwent 15-min on-column DNase digestion using an RNase-Free DNase Set (79254, Qiagen). cDNA was prepared with reverse transcriptase using an Omniscript RT kit (205113, Qiagen) following the standard protocol provided. RT-PCR was performed using EvaGreen (27490, Axon) kit with CFX96 Real Time System (BioRad). The standard two step-program was used: 95 °C for 10 min, then 40 cycles at 95 °C for 15 sec, 60 °C for 1 min. The expressions of *Gfap, Itgam*, *Pdgfra, Pdgfrb, Cxcl1, Cxcl10, Ccl2, Ccl5, Lcn2, Il6, Il1a, Il1b, Tnf* were measured. β-actin was performed as a control in all qPCR analysis. Primers were designed to work at approximately + 60°C and the specificity was assessed by melt curve analysis of each reaction indicating a single peak. Primer used in this study are listed in supplementary table S1. Relative expression of all targeted genes was determined using the ΔΔCt method with normalization to β-actin expression.

### Immunohistochemistry

Anesthetized mice were perfused with 1x PBS and 4% paraformaldehyde (PFA). Subsequently, the brain was dissected and post fixed in 4 % PFA overnight (4°C). Fixed brains were sliced into coronal sections of 35 μm thickness, in PBS, using a Leica VT1000S vibratome and collected in 48-well culture plates containing 1x PBS. Vibratome sections were incubated for 1 h in blocking buffer (5 % HS, 0.3 % Triton X in 1x PBS) at room temperature. Sections were incubated with primary antibodies, diluted in the blocking solution overnight at 4°C. The next day, sections were washed 3 times (10 min, 1x PBS) and incubated for 2 h with secondary antibodies and DAPI diluted in blocking buffer at room temperature in the dark. Subsequently, the sections were washed and mounted in Shandon ImmuMount (Thermo Scientific). Primary and secondary antibodies are listed in the key resources table, respectively.

### Image acquisition and analysis

Whole brain slices were scanned with the automatic slide scanner AxioScan.Z1 (Zeiss, Oberkochen) as described before(Huang et al., 2020). Cell counting from whole brain slices was manually performed using ZEN software (Zeiss, Jena). cFos intensity within Sox9^+^ and NeuN^+^ areas were automatically measured by using the machine learning function of the ZEN software. For the 3D reconstruction of microglia, confocal images were taken by LSM 710 and LSM780 confocal microscope (Zeiss, Oberkochen) with 1 μm intervals. Reconstruction of the microglial surface was performed using IMARIS (Version 9.6, Oxford Instruments) at the following settings: surface detail 0.700 μm (smooth); thresholding background subtraction (local contrast), diameter of largest sphere: 2.00. Next, the surface reconstruction was used as template for filament reconstruction with the following settings: detect new starting points: largest diameter 7.00 μm, seed points 0.300 μm; remove seed points around starting points: diameter of sphere regions: 15 μm. After filament reconstruction, individual data sets of Sholl analysis were exported into separate Excel files for further analysis. Image processing, three-dimensional reconstruction, and data analysis were performed in a blind manner regarding the experimental conditions.

### Ribosome immunoprecipitation (IP)

After perfusion with ice-cold HBSS, cortical regions samples were dissected from mouse brain and stored at − 80°C until use. Tissues were homogenized in ice-cold lysis buffer (50□mM Tris, pH 7.4, 100□mM KCl, 12□mM MgCl_2_, 1% NP-40, 1□mM DTT, 1x protease inhibitor, 200 units/ml RNasin (Promega) and 0.1□mg/ml cycloheximide (Sigma-Aldrich) in RNase-free deionized H_2_O) 10% w/v with homogenizer (Precellys 24, PeQlab). Then homogenates were centrifuged at 10,000□g at 4°C for 10□min to remove cell debris. Supernatants were collected and removed: 50 μl were used as input analysis. Anti-HA Ab (1:100, Covance) was added to the supernatant with slow rotation at 4°C. Meanwhile, Dynabeads Protein G (Thermo Fisher Scientific) were equilibrated to the lysis buffer by washing three times. After 4□h of incubation with HA Ab, 100 μl pre-equilibrated beads were added to each sample and incubated overnight at 4°C. After 10-12□h, Dynabeads were washed with high-salt buffer (50□mM Tris, 300□mM KCl, 12□mM MgCl_2_, 1% NP-40, 1□mM DTT, 1x protease inhibitor, 100 units/ml RNasin and 0.1□mg/ml cycloheximide in RNase-free deionized H_2_O) three times for 5 min at 4°C. At the end of the washing, beads were magnetized and 350□µl RLT buffer from RNeasy Micro Kit (74004, Qiagen) was added to the beads. RNA was extracted followed with manufacturer’s instructions.

### Next-generation RNA sequencing

The library was prepared and sequenced by Novogene using the following methods. Preliminary quality control was performed on 1% agarose gel electrophoresis to test RNA degradation and potential contamination. Sample purity and preliminary quantitation were measured using Bioanalyser 2100 (Agilent Technologies, USA), which was also used to check the RNA integrity and final quantitation. For library preparation, the mRNA present in the total RNA sample was isolated with magnetic beads of oligos d(T)25. This method is known as polyA-tailed mRNA enrichment. Subsequently, mRNA was randomly fragmented and cDNA synthesis proceeds using random hexamers and the reverse transcriptase enzyme. Once the synthesis of the first chain is finished, the second chain is synthesized with the addition of an Illumina buffer. Together with the presence of dNTPs, RNase H and polymerase I from E.coli, the second chain will be obtained by nick translation. The resulting products go through purification, end-repair, A-tailing and adapter ligation. Fragments of the appropriate size are enriched by PCR, where indexed P5 and P7 primers are introduced, and final products are purified. The library was checked with Qubit 2.0 and real-time PCR for quantification and bioanalyzer Agilent 2100 for size distribution detection. Quantified libraries were pooled and sequenced on Illumina Novaseq 6000 platform, according to effective library concentration and data amount.

The qualified libraries were sequenced Next Generation Sequencing (NGS) based on Illumina’s Sequencing Technology by Synthesis (SBS)-detection by fluorescence of the nucleotide added during the synthesis of the complementary chain - and in a parallelized and massive way. The Novaseq6000 sequencing system was used to sequence the libraries. The strategy is paired end 150bp (PE150).

### RNA-seq data processing

The RNA sequencing reads quality was assessed using FastQC (https://www.bioinformatics.babraham.ac.uk/projects/fastqc/). Reads were aligned to the GRCm38 Mus musculus genome using HISAT2 v2.0.5(Kim et al., 2019) with default parameters. Gene count matrix of each sample was generated by featureCounts v1.5.0-p3 (Liao et al., 2014). Downstream analysis was performed with the DEseq2 v1.20.0 (Love et al., 2014) package in R. Genes with normalized count below 10 were removed from downstream analysis. Significantly deregulated genes were identified by a false discovery rate lower than 0.05. Differential expression analysis between conditions was performed with the ‘DESeq’ function with default parameters. Log foldchange shrinkage was performed on the differential expression analysis result. Heatmaps of DEGs with p^adj^ value < 0.05 in any time point were visualized with pheatmap v1.0.12(Kolde, 2019). Selected gene set enrichment analysis of gene ontology (GO) and Kyoto Encyclopedia of Genes and Genomes (KEGG) pathway was performed with ClusterProfiler v3.8.1(Yu et al., 2012). Transcription factor prediction was performed using Metascape v3.5.20230501(Han et al., 2018; Zhou et al., 2019) with the differentially expressed genes identified by DEseq2.

### Cytokine expression analysis

After perfusion with ice-cold PBS, the cortex and striatum were dissected from coronal brain slices (1 mm) and stored at −80°C until further processing. For the cytokine array on brain homogenates, we used the Proteome Profiler Mouse Cytokine Array Kit (ARY006, R&D Systems) and Proteome Profiler Mouse XL Cytokine Array (ARY028, R&D Systems) following the manufacturer’s instructions. Briefly, brain tissue was homogenized in PBS containing 1x protease inhibitor cocktail (05892970001, Roche). Protein concentration was measured using the Bicinchoninic Acid (BCA) assay kit (Thermo Fisher Scientific). 200 μg protein lysis were used for each membrane. Membranes were blocked, incubated, and washed according to standard protocol. Four membranes were exposed to X-ray film (47410, Fuji) for 15 minutes at the same time. The intensity (pixel density) of each spot-on membrane was quantified using Image J software (National Institutes of Health, Bethesda), and corrected for background intensity and normalized to control.

### Acute brain slice preparation

Acute brain slice preparation was performed as previously described(Zhao et al., 2021a). Briefly, 11- to 13-week-old mice were anesthetized with isofluran (Abbvie, Ludwigshafen, Germany) and euthanized by cervical dislocation followed by decapitation. The brain was swiftly dissected and placed into ice-cooled, carbogen-saturated (5% CO_2_, 95% O_2_) cutting solution containing 87 mM NaCl, 3 mM KCl, 25 mM NaHCO_3_, 1.25 mM NaH_2_PO_4_, 3 mM MgCl_2_, 0.5 mM CaCl_2_, 75 mM sucrose, and 25 mM glucose. 300 μm-thick sections in correspondence to the primary somatosensory cortex (SSp-ctx, AP -1.6 ± 0.6) were cut with a vibratome VT1200S (Leica Biosystems, Wetzlar, DE) using a 0.12 mm/s cutting speed and a 1.9 mm cutting amplitude. The procedure between decapitation and incubation was performed as fast as possible and in no longer than 10 min for optimal quality of the slice preparation. Slices were incubated for 30 min on a custom-made nylon-basket submersed in artificial cerebral spinal fluid (ACSF) containing 126 mM NaCl, 3 mM KCl, 25 mM NaHCO_3_, 15 mM glucose, 1.2 mM NaH_2_PO_4_, 1 mM CaCl_2_, and 2 mM MgCl_2_ at 32°C. Subsequently, brain slices were taken out of the water bath and placed to RT with continuous oxygenation before use.

### Ca^2+^ imaging in hippocampal slices

For acute brain slices of 11- to 13-week-old GLAC^ct2/wt^xA1AR^wt/wt^xGCaMP3^fl/fl^ mice and GLAC^ct2/wt^xA1AR^fl/fl^xGCaMP3^fl/fl^ mice were prepared as described above. Individual slices were transferred to a self-made imaging chamber under a custom-made 2P-LSM and fixed by stainless steel rings with 1 mm-spaced nylon fibres (Harp Slice Grid HSG-5A; ALA Scientific Instruments Inc., Farmingdale, NY, USA). A constant flow of oxygenated perfusion solution continuously perfused the imaging chamber at a flow rate of 2–5 ml/min by a peristaltic pump LKB P-1 (Pharmacia LKB, Uppsala, SE). Squared field of views (FOVs, 170 x 170 μm) from the cortical layers II-III of the SSp-ctx were chosen at a depth ranging from 30 to 100 μm from the slice surface and displaying a uniform astroglial distribution. Focal CPA (0.1 mg/ml) and SR101 (0.1 mg/ml) application was performed by means of borosilicate glass pipettes (BF150-86-10, Sutter Instrument, Novato, CA, US) mounted on 3-axis micromanipulator units (LN Mini 25, Luigs & Neumann GmbH, Ratingen, DE) controlled by a SM5 Remote Control station (Luigs & Neumann GmbH, Ratingen, DE). Pipettes were pulled using a micropipette Puller P-97 (Sutter Instrument, Novato, CA, US) to obtain a 5 μm- opening tip (0.5-1 MΩ). Cortical astrocytic Ca^2+^ activity was recorded upon drug application. Analysis of Ca^2+^ imaging data was performed using the custom-made detection and analysis software MATLAB (MSparkles v. 1.8.18)(Stopper et al., 2023) as previously described (Rieder et al., 2022). Shortly, fluorescence fluctuations at basal Ca^2+^ concentrations (F_0_) were computed along the temporal axes of each individual pixel using a polynomial fitting in a least-squares sense. The range projection of ΔF/F0 was then used to identify local fluorescence maxima, serving as seed points for simultaneous, correlation-based growing of regions of activity (ROAs).

### Electrophysiology of brain slices (LTP)

LTP recordings from dorsal hippocampus were performed as previously described(Wang et al., 2016; Zhao et al., 2021b). Briefly, acute hippocampal brain slices were prepared as mentioned above. Afterwards, slices were transferred to the recording chamber and continuously perfused with oxygenated ACSF containing (in mM)1 MgCl2 and 2.5 CaCl2 at a flow rate of 2–5 mL/min. Field excitatory postsynaptic potentials (fEPSP) were recorded by a micropipette of 1–3 MΩ resistance filled with ACSF in CA1 of hippocampus by stimulating Schaffer collaterals of CA3 using stimulus isolator and a biopolar electrode (WPI). Picrotoxin (50 μM) was perfused in the bath to inhibit ionotropic γ-aminobutyric acid type A receptors (GABAARs). Stimulus duration was 200 μs, current injection was 30–80 μA. To evoke LTP, triple θ-burst stimulation (TBS3) was used. TBS consisted of 10 bursts (4 pulses each burst, 100 Hz) delivered at an interburst interval of 200 ms, and repeated once at 10 s. The stimulation intensity was adjusted to evoke ∼30–60% of the maximum response. Waveform analysis was performed by Igor pro 6.3.7.2. The statistical analysis was conducted in Graphpad Prism. All experiments were conducted at RT.

### Behavioral test

To evaluate the depression-like behavior following the LPS challenge, we evaluated open field test and sucrose preference test following the methods from previous studies with modifications (Liu et al., 2018; Fang et al., 2022). For the open field test, mice were placed at a random corner of the open field square (50□cm length□×□50□cm width ×□38□cm height) and positioned facing the wall. The movement of each mouse was recorded for 10 minutes. Open field test was performed before and after LPS challenge. Duration time in the center area (s), moved distance (cm) and speed (cm/s) were analyzed by EthoVision XT 11.5 (Noldus Technology). For the sucrose preference test, mice were single caged and habituated to the presence of two drinking bottles for 2 days. After the acclimatization, mice were presented to two drinking bottles: one containing 1% sucrose and the other tap water for 2 days in their cage. The positions of two drinking bottles were exchanged daily. Water and sucrose solution intake was measured daily by weighing the bottles. Sucrose preference was calculated as a percentage of the volume of sucrose intake over the total volume of fluid intake and averaged over the 2 days of testing. Sucrose preference test was performed before and after LPS challenge.

### *In vivo* activation of hM4Di

Three weeks after appropriate microinjection of AAV2/5-GFAP-hM4Di-mCherry or AAV2/5-GFAP-tdTomato into the striatum or cortex, CNO was administered two times to animals by intraperitoneal injection (2 mg/kg; dissolved in saline) at 2 hours and 4 hours post LPS challenge, respectively. 6 hours or 24 hours after LPS challenge, animals were used for behavior tests or immunohistochemistry.

### Statistics and reproducibility

The statistical analyses of all data were performed with GraphPad Prism 9.5.1 statistical software (GraphPad, San Diego). For all immunostainings, two randomly selected brain slices of each mouse were studied. In addition, for the analysis of Iba1, more than 8 ROIs per mouse was analyzed. For normally distributed dataset, unpaired t-tests, paired t-test (for studies of behavior), one-way ANOVA and two-way ANOVA were used (indicated in each figure legend), while the Kruskal-Wallis test was used for non-normally distributed datasets. P-values are indicated in the figures and legends. For the *in vivo* experiments, each data point represents the data obtained from a single mouse (except for electrophysiology). The total mouse numbers are indicated in the figure legends and table S2. For electrophysiology, each data point refers to a slice and the mouse number is indicated in the figure legends. Normal distributed datasets are shown as mean□±□SEM. else are shown as the median ± IQR are indicated as thick and thin dashed lines, respectively. *p-*value of ≤ 0.05 was considered statistically significant.

## References

Agnew-Svoboda W, Ubina T, Figueroa Z, Wong YC, Vizcarra EA, Roebini B, Wilson EH, Fiacco TA, Riccomagno MM (2022) A genetic tool for the longitudinal study of a subset of post-inflammatory reactive astrocytes. Cell Rep Methods 2:100276.

Benarroch E (2008) Adenosine and its receptors - Multiple modulatory functions and potential therapeutic targets for neurologic disease. Neurology 70:231–236.

Boison D (2008) Adenosine as a neuromodulator in neurological diseases. Current Opinion in Pharmacology 8:2–7.

Borea PA, Gessi S, Merighi S, Varani K (2016) Adenosine as a Multi-Signalling Guardian Angel in Human Diseases: When, Where and How Does it Exert its Protective Effects? Trends Pharmacol Sci 37:419–434.

Borea PA, Gessi S, Merighi S, Vincenzi F, Varani K (2017) Pathological overproduction: the bad side of adenosine. Br J Pharmacol 174:1945–1960.

Borst K, Dumas AA, Prinz M (2021) Microglia: Immune and non-immune functions. Immunity 54:2194–2208.

Bynoe MS, Viret C, Yan A, Kim DG (2015) Adenosine receptor signaling: a key to opening the blood-brain door. Fluids Barriers CNS 12:20.

Carman AJ, Mills JH, Krenz A, Kim DG, Bynoe MS (2011) Adenosine receptor signaling modulates permeability of the blood-brain barrier. J Neurosci 31:13272–13280.

Chiu G, Freund G (2014) Modulation of neuroimmunity by adenosine and its receptors: Metabolism to mental illness. Metabolism-Clinical and Experimental 63:1491–1498.

Chiu GS, Darmody PT, Walsh JP, Moon ML, Kwakwa KA, Bray JK, McCusker RH, Freund GG (2014) Adenosine through the A2A adenosine receptor increases IL-1β in the brain contributing to anxiety. Brain Behav Immun 41:218–231.

Cupido A, Catalin B, Steffens H, Kirchhoff F (2014) Surgical procedures to study microglial motility in the brain and in the spinal cord by in vivo two-photon laser-scanning microscopy. Laser scanning microscopy and quantitative image analysis of neuronal tissue:37–50.

Dantzer R, O’Connor JC, Freund GG, Johnson RW, Kelley KW (2008) From inflammation to sickness and depression: when the immune system subjugates the brain. Nat Rev Neurosci 9:46–56.

Duan L, Zhang XD, Miao WY, Sun YJ, Xiong G, Wu Q, Li G, Yang P, Yu H, Li H, Wang Y, Zhang M, Hu LY, Tong X, Zhou WH, Yu X (2018) PDGFRβ Cells Rapidly Relay Inflammatory Signal from the Circulatory System to Neurons via Chemokine CCL2. Neuron 100:183–200.e188.

Fang L-P, Zhao N, Caudal LC, Chang H-F, Zhao R, Lin C-H, Hainz N, Meier C, Bettler B, Huang W (2022) Impaired bidirectional communication between interneurons and oligodendrocyte precursor cells affects social cognitive behavior. Nature Communications 13:1394.

Fritz M et al. (2016) Prostaglandin-dependent modulation of dopaminergic neurotransmission elicits inflammation-induced aversion in mice. J Clin Invest 126:695–705.

Han H, Cho JW, Lee S, Yun A, Kim H, Bae D, Yang S, Kim CY, Lee M, Kim E, Lee S, Kang B, Jeong D, Kim Y, Jeon HN, Jung H, Nam S, Chung M, Kim JH, Lee I (2018) TRRUST v2: an expanded reference database of human and mouse transcriptional regulatory interactions. Nucleic Acids Res 46:D380–d386.

Han RT, Kim RD, Molofsky AV, Liddelow SA (2021) Astrocyte-immune cell interactions in physiology and pathology. Immunity 54:211–224.

Haruwaka K, Ikegami A, Tachibana Y, Ohno N, Konishi H, Hashimoto A, Matsumoto M, Kato D, Ono R, Kiyama H, Moorhouse AJ, Nabekura J, Wake H (2019) Dual microglia effects on blood brain barrier permeability induced by systemic inflammation. Nat Commun 10:5816.

Hasel P, Rose IVL, Sadick JS, Kim RD, Liddelow SA (2021a) Neuroinflammatory astrocyte subtypes in the mouse brain. Nat Neurosci 24:1475–1487.

Hasel P, Rose IV, Sadick JS, Kim RD, Liddelow SA (2021b) Neuroinflammatory astrocyte subtypes in the mouse brain. Nat Neurosci 24:1475–1487.

He Q, Wang J, Hu H (2019) Illuminating the Activated Brain: Emerging Activity-Dependent Tools to Capture and Control Functional Neural Circuits. Neurosci Bull 35:369–377.

Huang W, Bai X, Meyer E, Scheller A (2020) Acute brain injuries trigger microglia as an additional source of the proteoglycan NG2. Acta Neuropathol Commun 8:146.

Huang X, Guo M, Zhang Y, Xie J, Huang R, Zuo Z, Saw PE, Cao M (2023) Microglial IL-1RA ameliorates brain injury after ischemic stroke by inhibiting astrocytic CXCL1-mediated neutrophil recruitment and microvessel occlusion. Glia 71:1607–1625.

Hösli L, Zuend M, Bredell G, Zanker HS, Porto de Oliveira CE, Saab AS, Weber B (2022) Direct vascular contact is a hallmark of cerebral astrocytes. Cell Rep 39:110599.

Izumi Y, Cashikar AG, Krishnan K, Paul SM, Covey DF, Mennerick SJ, Zorumski CF (2021) A Proinflammatory Stimulus Disrupts Hippocampal Plasticity and Learning via Microglial Activation and 25-Hydroxycholesterol. J Neurosci 41:10054–10064.

Kang SS, Ren Y, Liu CC, Kurti A, Baker KE, Bu G, Asmann Y, Fryer JD (2018) Lipocalin-2 protects the brain during inflammatory conditions. Mol Psychiatry 23:344–350.

Kim D, Paggi JM, Park C, Bennett C, Salzberg SL (2019) Graph-based genome alignment and genotyping with HISAT2 and HISAT-genotype. Nat Biotechnol 37:907-+.

Kislin M, Mugantseva E, Molotkov D, Kulesskaya N, Khirug S, Kirilkin I, Pryazhnikov E, Kolikova J, Toptunov D, Yuryev M (2014) Flat-floored air-lifted platform: a new method for combining behavior with microscopy or electrophysiology on awake freely moving rodents. JoVE (Journal of Visualized Experiments):e51869.

Klawonn AM, Fritz M, Castany S, Pignatelli M, Canal C, Similä F, Tejeda HA, Levinsson J, Jaarola M, Jakobsson J, Hidalgo J, Heilig M, Bonci A, Engblom D (2021) Microglial activation elicits a negative affective state through prostaglandin-mediated modulation of striatal neurons. Immunity 54:225–234.e226.

Kodali MC, Chen H, Liao FF (2021) Temporal unsnarling of brain’s acute neuroinflammatory transcriptional profiles reveals panendothelitis as the earliest event preceding microgliosis. Mol Psychiatry 26:3905–3919.

Kolde R (2019) Pheatmap: Pretty Heatmaps, R Package Version 1.0. 12. 2019. In.

Latini S, Pedata F (2001) Adenosine in the central nervous system: release mechanisms and extracellular concentrations. J Neurochem 79:463–484.

Liao Y, Smyth GK, Shi W (2014) featureCounts: an efficient general purpose program for assigning sequence reads to genomic features. Bioinformatics 30:923–930.

Liddelow SA et al. (2017) Neurotoxic reactive astrocytes are induced by activated microglia. Nature 541:481–487.

Linnerbauer M, Wheeler MA, Quintana FJ (2020) Astrocyte Crosstalk in CNS Inflammation. Neuron 108:608–622.

Liu MY, Yin CY, Zhu LJ, Zhu XH, Xu C, Luo CX, Chen HS, Zhu DY, Zhou QG (2018) Sucrose preference test for measurement of stress-induced anhedonia in mice. Nat Protoc 13:1686–1698.

Liu Q, Guo Q, Fang LP, Yao H, Scheller A, Kirchhoff F, Huang W (2023) Specific detection and deletion of the sigma-1 receptor widely expressed in neurons and glial cells in vivo. J Neurochem 164:764–785.

Love MI, Huber W, Anders S (2014) Moderated estimation of fold change and dispersion for RNA-seq data with DESeq2. Genome Biol 15.

Manabe T, Heneka MT (2022) Cerebral dysfunctions caused by sepsis during ageing. Nat Rev Immunol 22:444–458.

Martin C, Leone M, Viviand X, Ayem ML, Guieu R (2000) High adenosine plasma concentration as a prognostic index for outcome in patients with septic shock. Crit Care Med 28:3198–3202.

Mazeraud A, Righy C, Bouchereau E, Benghanem S, Bozza FA, Sharshar T (2020) Septic-Associated Encephalopathy: a Comprehensive Review. Neurotherapeutics 17:392–403.

Meng F, Guo Z, Hu Y, Mai W, Zhang Z, Zhang B, Ge Q, Lou H, Guo F, Chen J, Duan S, Gao Z (2019) CD73-derived adenosine controls inflammation and neurodegeneration by modulating dopamine signalling. Brain 142:700–718.

Mikitsh JL, Chacko AM (2014) Pathways for small molecule delivery to the central nervous system across the blood-brain barrier. Perspect Medicin Chem 6:11–24.

Mills JH, Thompson LF, Mueller C, Waickman AT, Jalkanen S, Niemela J, Airas L, Bynoe MS (2008) CD73 is required for efficient entry of lymphocytes into the central nervous system during experimental autoimmune encephalomyelitis. Proc Natl Acad Sci U S A 105:9325–9330.

Mori T, Tanaka K, Buffo A, Wurst W, Kuhn R, Gotz M (2006) Inducible gene deletion in astroglia and radial glia - A valuable tool for functional and lineage analysis. Glia 54:21–34.

Motori E, Puyal J, Toni N, Ghanem A, Angeloni C, Malaguti M, Cantelli-Forti G, Berninger B, Conzelmann KK, Götz M, Winklhofer KF, Hrelia S, Bergami M (2013) Inflammation-induced alteration of astrocyte mitochondrial dynamics requires autophagy for mitochondrial network maintenance. Cell Metab 18:844–859.

Nagai J, Rajbhandari AK, Gangwani MR, Hachisuka A, Coppola G, Masmanidis SC, Fanselow MS, Khakh BS (2019a) Hyperactivity with disrupted attention by activation of an astrocyte synaptogenic cue. Cell 177:1280–1292. e1220.

Nagai J, Rajbhandari AK, Gangwani MR, Hachisuka A, Coppola G, Masmanidis SC, Fanselow MS, Khakh BS (2019b) Hyperactivity with Disrupted Attention by Activation of an Astrocyte Synaptogenic Cue. Cell 177:1280–1292.e1220.

Patani R, Hardingham GE, Liddelow SA (2023) Functional roles of reactive astrocytes in neuroinflammation and neurodegeneration. Nat Rev Neurol 19:395–409.

Paukert M, Agarwal A, Cha J, Doze VA, Kang JU, Bergles DE (2014) Norepinephrine controls astroglial responsiveness to local circuit activity. Neuron 82:1263–1270.

Peng W, Wu Z, Song K, Zhang S, Li Y, Xu M (2020) Regulation of sleep homeostasis mediator adenosine by basal forebrain glutamatergic neurons. Science 369.

Ramakers BP, Riksen NP, van der Hoeven JG, Smits P, Pickkers P (2011a) Modulation of innate immunity by adenosine receptor stimulation. Shock 36:208–215.

Ramakers BP, Riksen NP, van den Broek P, Franke B, Peters WH, van der Hoeven JG, Smits P, Pickkers P (2011b) Circulating adenosine increases during human experimental endotoxemia but blockade of its receptor does not influence the immune response and subsequent organ injury. Crit Care 15:R3.

Rieder P, Gobbo D, Stopper G, Welle A, Damo E, Kirchhoff F, Scheller A (2022) Astrocytes and Microglia Exhibit Cell-Specific Ca. Front Mol Neurosci 15:840948.

Rummel C, Inoue W, Poole S, Luheshi GN (2010) Leptin regulates leukocyte recruitment into the brain following systemic LPS-induced inflammation. Mol Psychiatry 15:523–534.

Sanz E, Yang L, Su T, Morris DR, McKnight GS, Amieux PS (2009) Cell-type-specific isolation of ribosome-associated mRNA from complex tissues. Proc Natl Acad Sci U S A 106:13939–13944.

Scammell TE, Arrigoni E, Thompson MA, Ronan PJ, Saper CB, Greene RW (2003) Focal deletion of the adenosine A1 receptor in adult mice using an adeno-associated viral vector. J Neurosci 23:5762–5770.

Shemer A, Scheyltjens I, Frumer GR, Kim JS, Grozovski J, Ayanaw S, Dassa B, Van Hove H, Chappell-Maor L, Boura-Halfon S, Leshkowitz D, Mueller W, Maggio N, Movahedi K, Jung S (2020) Interleukin-10 Prevents Pathological Microglia Hyperactivation following Peripheral Endotoxin Challenge. Immunity 53:1033–1049.e1037.

Sofroniew MV (2020) Astrocyte Reactivity: Subtypes, States, and Functions in CNS Innate Immunity. Trends Immunol 41:758–770.

Stopper G, Caudal LC, Rieder P, Gobbo D, Stopper L, Felix L, Everaerts K, Bai X, Rose CR, Scheller A, Kirchhoff F (2023) Novel algorithms for improved detection and analysis of fluorescent signal fluctuations. Pflugers Arch.

Sumi Y, Woehrle T, Chen Y, Bao Y, Li X, Yao Y, Inoue Y, Tanaka H, Junger WG (2014) Plasma ATP is required for neutrophil activation in a mouse sepsis model. Shock 42:142–147.

van der Poll T, van de Veerdonk FL, Scicluna BP, Netea MG (2017) The immunopathology of sepsis and potential therapeutic targets. Nat Rev Immunol 17:407–420.

Wang H, Ardiles AO, Yang S, Tran T, Posada-Duque R, Valdivia G, Baek M, Chuang YA, Palacios AG, Gallagher M, Worley P, Kirkwood A (2016) Metabotropic Glutamate Receptors Induce a Form of LTP Controlled by Translation and Arc Signaling in the Hippocampus. J Neurosci 36:1723–1729.

Welsh TG, Kucenas S (2018) Purinergic signaling in oligodendrocyte development and function. J Neurochem 145:6–18.

Wu KC, Lee CY, Chern Y, Lin CJ (2021) Amelioration of lipopolysaccharide-induced memory impairment in equilibrative nucleoside transporter-2 knockout mice is accompanied by the changes in glutamatergic pathways. Brain Behav Immun 96:187–199.

Wu Z, He K, Chen Y, Li H, Pan S, Li B, Liu T, Xi F, Deng F, Wang H, Du J, Jing M, Li Y (2022) A sensitive GRAB sensor for detecting extracellular ATP in vitro and in vivo. Neuron 110:770–782.e775.

Ximerakis M, Lipnick SL, Innes BT, Simmons SK, Adiconis X, Dionne D, Mayweather BA, Nguyen L, Niziolek Z, Ozek C, Butty VL, Isserlin R, Buchanan SM, Levine SS, Regev A, Bader GD, Levin JZ, Rubin LL (2019) Single-cell transcriptomic profiling of the aging mouse brain. Nat Neurosci 22:1696–1708.

Yu G, Wang L-G, Han Y, He Q-Y (2012) clusterProfiler: an R package for comparing biological themes among gene clusters. Omics: a journal of integrative biology 16:284–287.

Zhang Y, Chen K, Sloan S, Bennett M, Scholze A, O’Keeffe S, Phatnani H, Guarnieri P, Caneda C, Ruderisch N, Deng S, Liddelow S, Zhang C, Daneman R, Maniatis T, Barres B, Wu J (2014) An RNA-Sequencing Transcriptome and Splicing Database of Glia, Neurons, and Vascular Cells of the Cerebral Cortex. Journal of Neuroscience 34:11929–11947.

Zhao N, Huang W, Cãtãlin B, Scheller A, Kirchhoff F (2021a) L-Type Ca2+ channels of NG2 glia determine proliferation and NMDA receptor-dependent plasticity. Frontiers in Cell and Developmental Biology 9:759477.

Zhao N, Huang W, Cãtãlin B, Scheller A, Kirchhoff F (2021b) L-Type Ca(2+) Channels of NG2 Glia Determine Proliferation and NMDA Receptor-Dependent Plasticity. Front Cell Dev Biol 9:759477.

Zhao Z, Shang X, Chen Y, Zheng Y, Huang W, Jiang H, Lv Q, Kong D, Jiang Y, Liu P (2020) Bacteria elevate extracellular adenosine to exploit host signaling for blood-brain barrier disruption. Virulence 11:980–994.

Zhou Y, Zhou B, Pache L, Chang M, Khodabakhshi AH, Tanaseichuk O, Benner C, Chanda SK (2019) Metascape provides a biologist-oriented resource for the analysis of systems-level datasets. Nature communications 10:1523.

