## Supplementary Figures 1-6 for "Adenosine triggers early astrocyte reactivity that provokes microglial activation and drives the pathogenesis of sepsis-associated encephalopathy"

Xianshu Bai: 0000-0002-4758-1645

Anja Scheller: 0000-0001-8955-2634

Frank Kirchhoff: 0000-0002-2324-2761

Wenhui Huang: 0000-0001-9865-0375

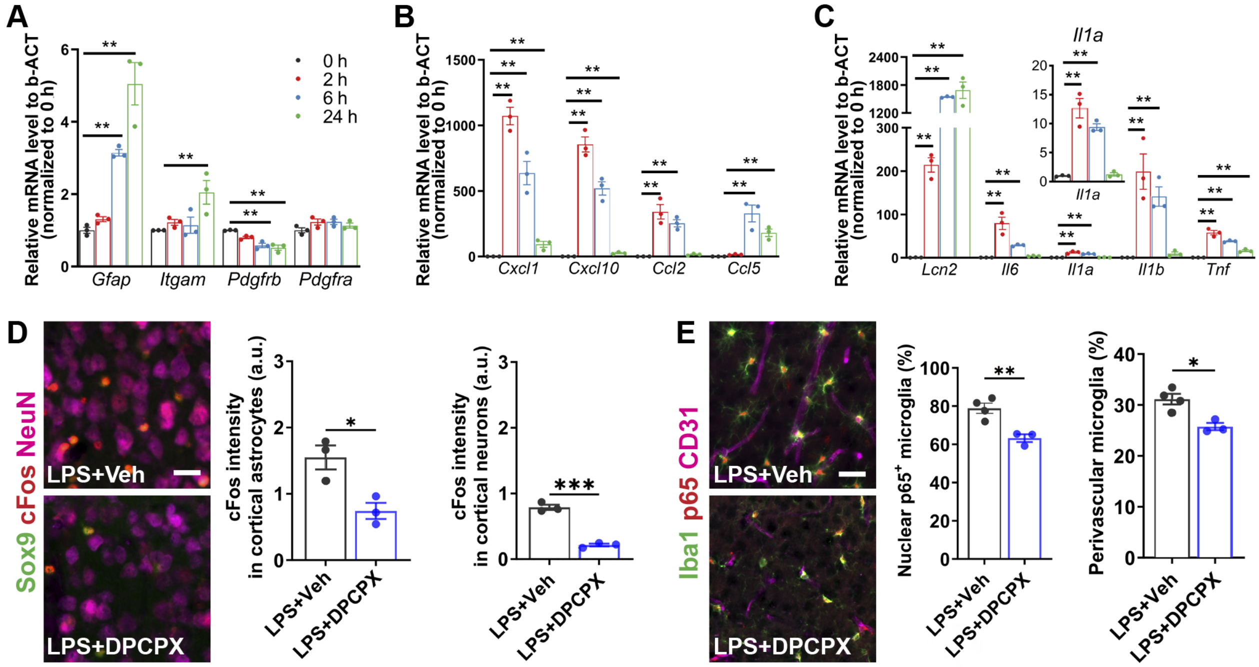


**Figure S1. Pharmacological inhibition of A1AR alleviates systemic inflammation-induced cytokine expression and glia activation; related to Figure 1 and 2.**

**(A)** Changes of marker genes for astrocytes (*Gfap*), microglia (*Itgam*), pericytes (*Pdgfrb*), and OPCs (*Pdgfra*) after PBS/LPS injection (n = 3 mice per group).

**(B, C)** Expression levels of several chemokines **(B)** and proinflammatory cytokines **(C)** after PBS/LPS injection (n = 3 mice per group).

**(D)** Representative images of immunoreactivity of c-Fos in Sox9$^{+}$ astrocytes and NeuN+ neurons in the mouse cortex upon LPS and A1AR antagonist (DPCPX) injection (left). c-Fos expression in astrocytes and neurons was reduced by DPCPX (right) (n = 3 mice per group).

**(E)** Representative images of immunolabeled nuclear p65$^{+}$ microglia and CD31$^{+}$ blood vessels upon LPS and A1AR antagonist (DPCPX) injection (left). Nuclear p65$^{+}$ microglia and perivascular microglia were reduced by DPCPX (right) (n = 4 mice in LPS+Veh group, n = 3 mice in LPS+DPCPX group).

Summary data are presented as the mean ± SEM. Statistical significance in (**A-C**) were assessed by two-way ANOVA; statistical significance in (**D, E**) was assessed by unpaired Student's t test, $^{*}P < 0.05$, $^{**}P < 0.01$, $^{***}P < 0.001$.

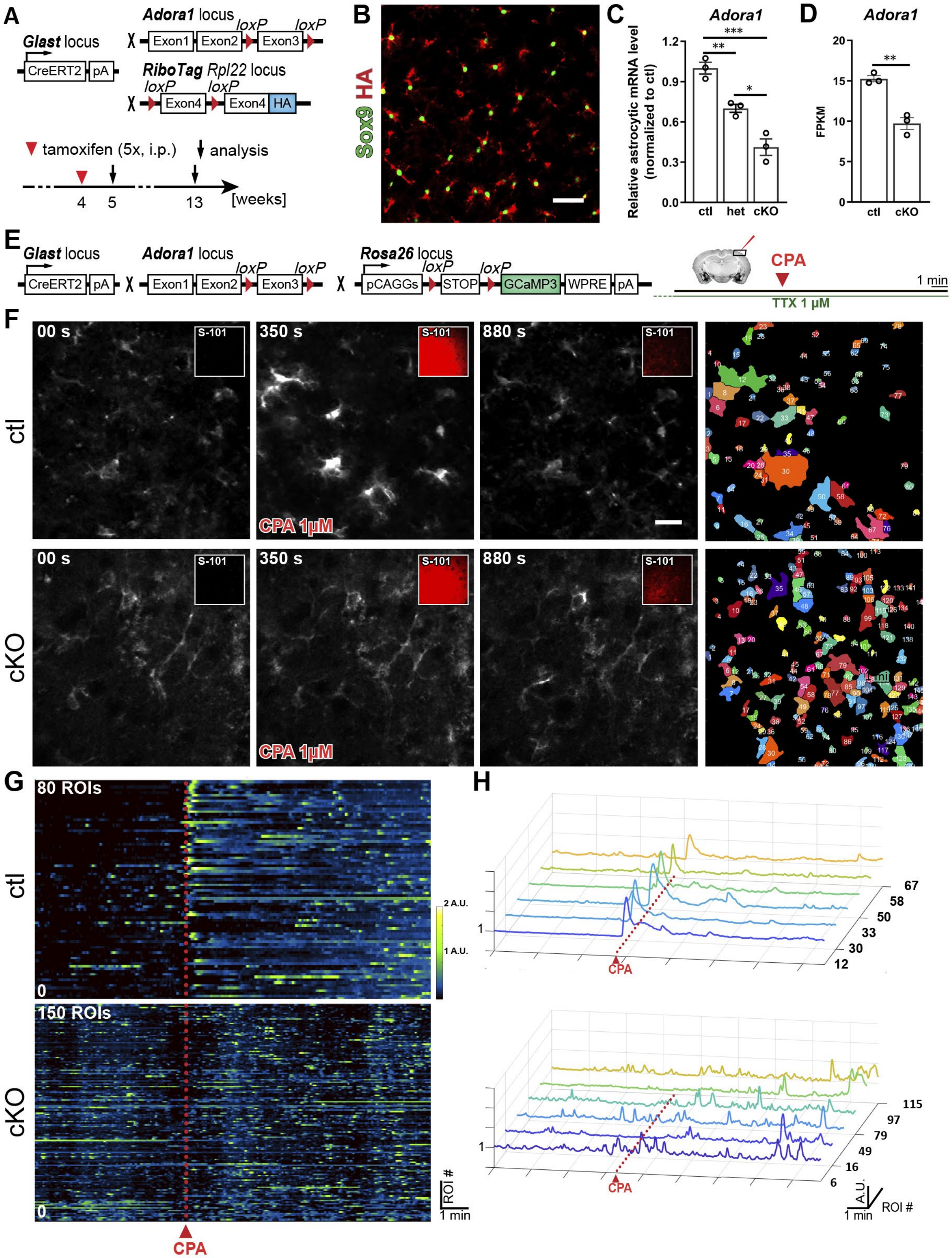


### Figure S2. Generation and validation of astrocyte-specific A1AR deficient mice; related to Figure 2 and 3.

**(A)** Schematic illustration of mouse breeding for astrocyte-specific A1AR deficient mice (cKO) and experiment plan. GLAST-CreERT2 (GLAC) mice were crossed to floxed A1AR mice. RiboTag mice were also introduced to the breeding for specifically and directly purify translated mRNA from astrocytes without sorting cells.

**(B)** Representative image of RiboTag expression (indicated by HA-tag) in $Sox9^+$ astrocytes. Scale bar = 50 µm.

**(C)** *Adora1* expression in astrocytes was reduced in $A1AR^{fl/wt}$ (het) and cKO mice one week after tamoxifen injection by using qPCR (n = 3 mice per group).

**(D)** *Adora1* expression in astrocytes was reduced in cKO mice 9 weeks after tamoxifen injection by using RNA-Seq (n = 3 mice per group).

**(E)** Schematic illustration of mouse breeding for $Ca^{2+}$ imaging and experiment plan. $GLACxA1AR^{fl/fl}$ mice were crossed to Rosa26-GCaMP3 mice. $GLACxA1AR^{fl/fl}$xRosa26-GCaMP3 mice were treated with tamoxifen at 4 weeks and used for *ex vivo* $Ca^{2+}$ imaging at 13 weeks of age. Coronal brain slices were incubated in TTX (tetrodotoxin). During recording the A1AR agonist CPA (1 µM) was applied focally. Sulforhodamine 101 (SR101, 4 µg/ml) was mixed with CPA to indicate the drug application.

**(F)** Images showing the change of $Ca^{2+}$ activity during the recording in ctl and $GLACxA1AR^{fl/fl}$ mice. Notably, CPA application evoked high $Ca^{2+}$ increase in ctl mice which was not observed in cKO mice, functionally confirming the deletion of A1ARs in astrocytes. The rightmost images show automatically detected regions of interests (ROIs) with dynamic $Ca^{2+}$ activities by a custom-made tool MSparkles.

**(G)** Heatmap plot showing amplitude and duration of spontaneous $Ca^{2+}$ events detected from all ROIs in **(F)**.

**(H)** Six ROIs in **(F)** were selected to show the characteristics of $Ca^{2+}$ events.

Summary data are presented as the mean ± SEM. Statistical significance in **(C)** was assessed using a one-way ANOVA; statistical significance in (**D**) was assessed using unpaired Student's t test, *P < 0.05, **P < 0.01, ***P < 0.001.

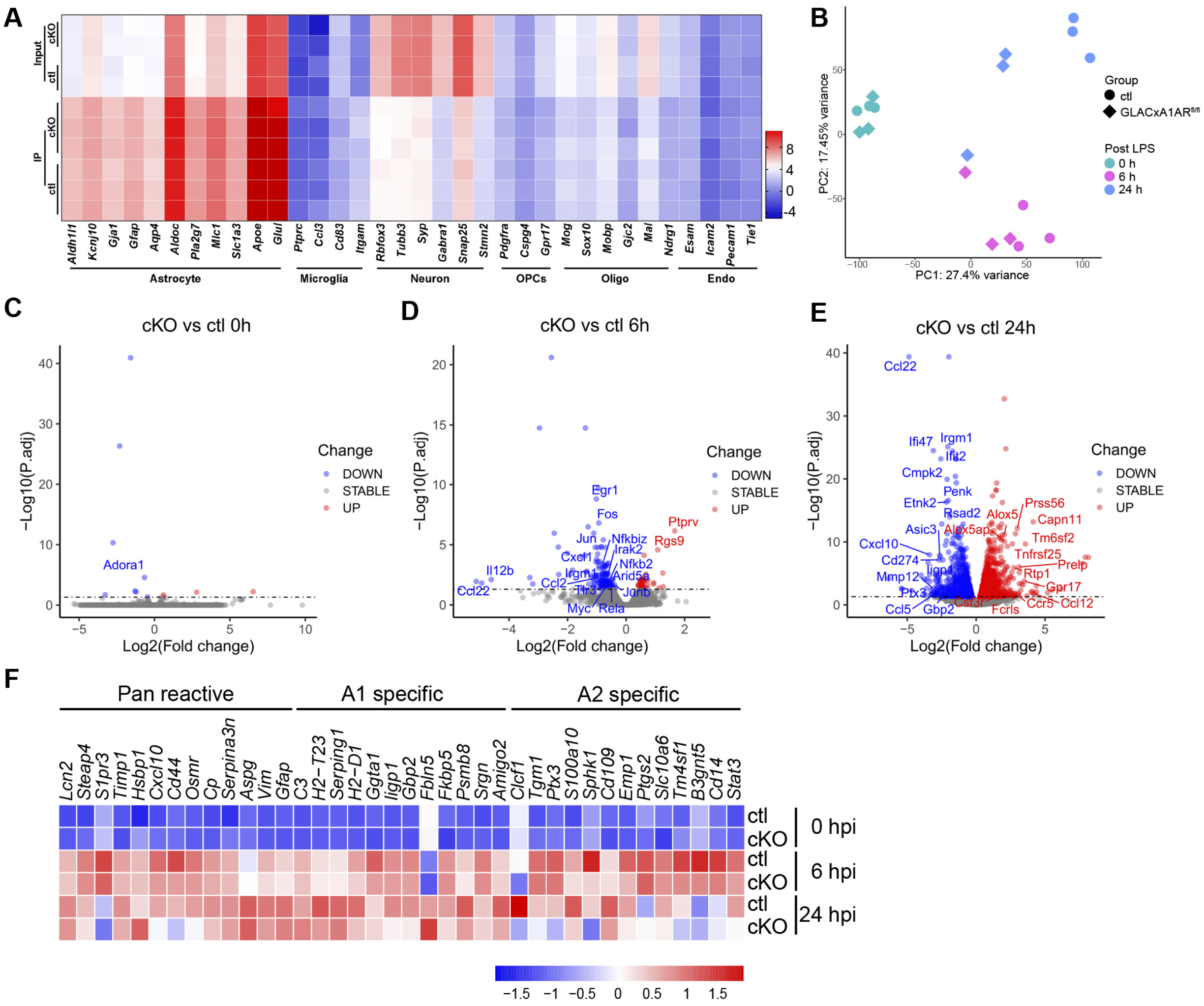


**Figure S3. A1AR-deficient astrocytes are less reactive to the peripheral LPS challenge; related to Figure 3.**

**(A)** Heatmap of cell type-specific marker gene expression showed immunoprecipitation (IP) of RiboTag enriched astrocyte-specific genes.

**(B)** PCA (principal component analysis) of the RNA-seq dataset (n = 3 mice per group).

**(C-E)** Volcano plots showing gene expression changes in cKO group compared to ctl group at 0 hpi **(C)**, 6 hpi **(D)**, 24 hpi **(E)**.

**(F)** Heatmap of marker genes for A1/A2 astrocytes.

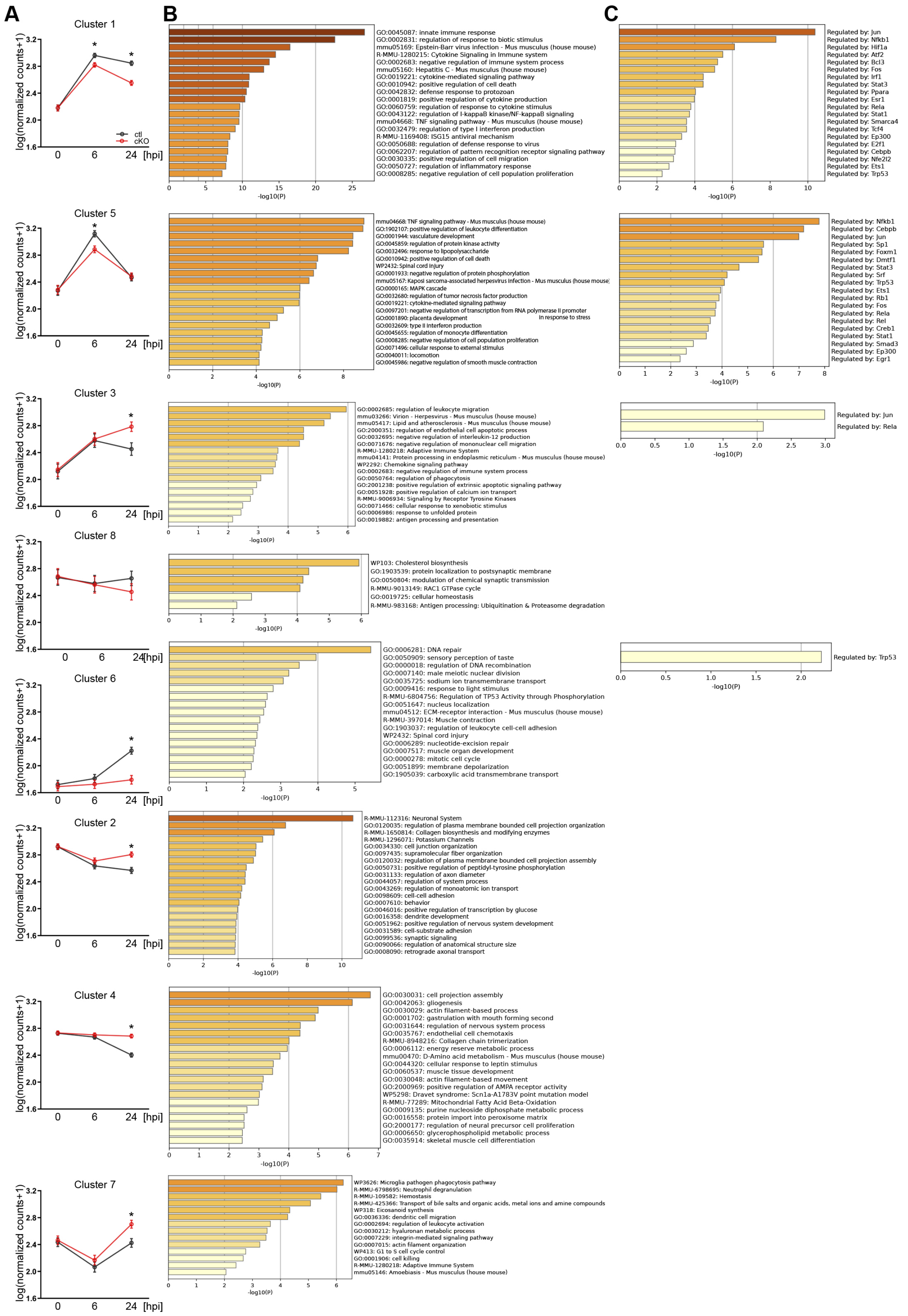

A
B
C
Cluster 1
log(normalized counts+1)
ctl
cKO
0
6
24 [hpi]
GO:0045087: innate immune response
GO:0002831: regulation of response to biotic stimulus
mmu05169: Epstein-Barr virus infection - Mus musculus (house mouse)
R-MMU-1280215: Cytokine Signaling in Immune system
GO:0002683: negative regulation of immune system process
mmu05160: Hepatitis C - Mus musculus (house mouse)
GO:0019221: cytokine-mediated signaling pathway
GO:0010942: positive regulation of cell death
GO:0042832: defense response to protozoan
GO:0001819: positive regulation of cytokine production
GO:0060759: regulation of response to cytokine stimulus
GO:0043122: regulation of I-kappaB kinase/NF-kappaB signaling
mmu04668: TNF signaling pathway - Mus musculus (house mouse)
GO:0032479: regulation of type I interferon production
R-MMU-1169408: ISG15 antiviral mechanism
GO:0050688: regulation of defense response to virus
GO:0062207: regulation of pattern recognition receptor signaling pathway
GO:0030335: positive regulation of cell migration
GO:0050727: regulation of inflammatory response
GO:0008285: negative regulation of cell population proliferation
-log10(P)
Regulated by: Jun
Regulated by: Nfkb1
Regulated by: Hif1a
Regulated by: Atf2
Regulated by: Bcl3
Regulated by: Fos
Regulated by: Irf1
Regulated by: Stat3
Regulated by: Ppara
Regulated by: Esr1
Regulated by: Rela
Regulated by: Stat1
Regulated by: Smarca4
Regulated by: Tcf4
Regulated by: Ep300
Regulated by: E2f1
Regulated by: Cebpb
Regulated by: Nfe2l2
Regulated by: Ets1
Regulated by: Trp53
Cluster 5
mmu04668: TNF signaling pathway - Mus musculus (house mouse)
GO:1902107: positive regulation of leukocyte differentiation
GO:0001944: vasculature development
GO:0045859: regulation of protein kinase activity
GO:0032496: response to lipopolysaccharide
GO:0010942: positive regulation of cell death
WP2432: Spinal cord injury
GO:0001933: negative regulation of protein phosphorylation
mmu05167: Kaposi sarcoma-associated herpesvirus infection - Mus musculus (house mouse)
GO:0000165: MAPK cascade
GO:0032680: regulation of tumor necrosis factor production
GO:0019221: cytokine-mediated signaling pathway
GO:0097201: negative regulation of transcription from RNA polymerase II promoter in response to stress
GO:0001890: placenta development
GO:0032609: type II interferon production
GO:0045655: regulation of monocyte differentiation
GO:0008285: negative regulation of cell population proliferation
GO:0071496: cellular response to external stimulus
GO:0040011: locomotion
GO:0045986: negative regulation of smooth muscle contraction
Regulated by: Nfkb1
Regulated by: Cebpb
Regulated by: Jun
Regulated by: Sp1
Regulated by: Foxm1
Regulated by: Dmtf1
Regulated by: Stat3
Regulated by: Srf
Regulated by: Trp53
Regulated by: Ets1
Regulated by: Rb1
Regulated by: Fos
Regulated by: Rela
Regulated by: Rel
Regulated by: Creb1
Regulated by: Stat1
Regulated by: Smad3
Regulated by: Ep300
Regulated by: Egr1
Cluster 3
GO:0002685: regulation of leukocyte migration
mmu03266: Virion - Herpesvirus - Mus musculus (house mouse)
mmu05417: Lipid and atherosclerosis - Mus musculus (house mouse)
GO:2000351: regulation of endothelial cell apoptotic process
GO:0032695: negative regulation of interleukin-12 production
GO:0071676: negative regulation of mononuclear cell migration
R-MMU-1280218: Adaptive Immune System
mmu04141: Protein processing in endoplasmic reticulum - Mus musculus (house mouse)
WP2292: Chemokine signaling pathway
GO:0002683: negative regulation of immune system process
GO:0050764: regulation of phagocytosis
GO:2001238: positive regulation of extrinsic apoptotic signaling pathway
GO:0051928: positive regulation of calcium ion transport
R-MMU-9006934: Signaling by Receptor Tyrosine Kinases
GO:0071466: cellular response to xenobiotic stimulus
GO:0006986: response to unfolded protein
GO:0019882: antigen processing and presentation
Regulated by: Jun
Regulated by: Rela
Cluster 8
WP103: Cholesterol biosynthesis
GO:1903539: protein localization to postsynaptic membrane
GO:0050804: modulation of chemical synaptic transmission
R-MMU-9013149: RAC1 GTPase cycle
GO:0019725: cellular homeostasis
R-MMU-983168: Antigen processing: Ubiquitination & Proteasome degradation
Cluster 6
GO:0006281: DNA repair
GO:0050909: sensory perception of taste
GO:0000018: regulation of DNA recombination
GO:0007140: male meiotic nuclear division
GO:0035725: sodium ion transmembrane transport
GO:0009416: response to light stimulus
R-MMU-6804756: Regulation of TP53 Activity through Phosphorylation
GO:0051647: nucleus localization
mmu04512: ECM-receptor interaction - Mus musculus (house mouse)
R-MMU-397014: Muscle contraction
GO:1903037: regulation of leukocyte cell-cell adhesion
WP2432: Spinal cord injury
GO:0006289: nucleotide-excision repair
GO:0007517: muscle organ development
GO:0000278: mitotic cell cycle
GO:0051899: membrane depolarization
GO:1905039: carboxylic acid transmembrane transport
Regulated by: Trp53
Cluster 2
R-MMU-112316: Neuronal System
GO:0120035: regulation of plasma membrane bounded cell projection organization
R-MMU-1650814: Collagen biosynthesis and modifying enzymes
R-MMU-1296071: Potassium Channels
GO:0034330: cell junction organization
GO:0097435: supramolecular fiber organization
GO:0120032: regulation of plasma membrane bounded cell projection assembly
GO:0050731: positive regulation of peptidyl-tyrosine phosphorylation
GO:0031133: regulation of axon diameter
GO:0044057: regulation of system process
GO:0043269: regulation of monoatomic ion transport
GO:0098609: cell-cell adhesion
GO:0007610: behavior
GO:0046016: positive regulation of transcription by glucose
GO:0016358: dendrite development
GO:0051962: positive regulation of nervous system development
GO:0031589: cell-substrate adhesion
GO:0099536: synaptic signaling
GO:0090066: regulation of anatomical structure size
GO:0008090: retrograde axonal transport
Cluster 4
GO:0030031: cell projection assembly
GO:0042063: gliogenesis
GO:0030029: actin filament-based process
GO:0001702: gastrulation with mouth forming second
GO:0031644: regulation of nervous system process
GO:0035767: endothelial cell chemotaxis
R-MMU-8948216: Collagen chain trimerization
GO:0006112: energy reserve metabolic process
mmu00470: D-Amino acid metabolism - Mus musculus (house mouse)
GO:0044320: cellular response to leptin stimulus
GO:0060537: muscle tissue development
GO:0030048: actin filament-based movement
GO:2000969: positive regulation of AMPA receptor activity
WP5298: Dravet syndrome: Scn1a-A1783V point mutation model
R-MMU-77289: Mitochondrial Fatty Acid Beta-Oxidation
GO:0009135: purine nucleoside diphosphate metabolic process
GO:0016558: protein import into peroxisome matrix
GO:2000177: regulation of neural precursor cell proliferation
GO:0006650: glycerophospholipid metabolic process
GO:0035914: skeletal muscle cell differentiation
Cluster 7
WP3626: Microglia pathogen phagocytosis pathway
R-MMU-6798695: Neutrophil degranulation
R-MMU-109582: Hemostasis
R-MMU-425366: Transport of bile salts and organic acids, metal ions and amine compounds
WP318: Eicosanoid synthesis
GO:0036336: dendritic cell migration
GO:0002694: regulation of leukocyte activation
GO:0030212: hyaluronan metabolic process
GO:0007229: integrin-mediated signaling pathway
GO:0007015: actin filament organization
WP413: G1 to S cell cycle control
GO:0001906: cell killing
R-MMU-1280218: Adaptive Immune System
mmu05146: Amoebiasis - Mus musculus (house mouse)

**Figure S4. Further transcriptomic data analysis reveals A1AR-deficient astrocytes are less reactive to the peripheral LPS challenge; related to Figure 3.**

**(A)** Mean profile representation of the temporal gene expression pattern for each cluster in Figure 3D. Data points correspond to 0 hpi, 6 hpi, 24 hpi.

**(B)** Metascape pathways for each cluster in Figure 3D generated by Metascape analysis.

**(C)** List of prediction of transcription regulators following expression pattern of sub-clusters in Figure 3D.

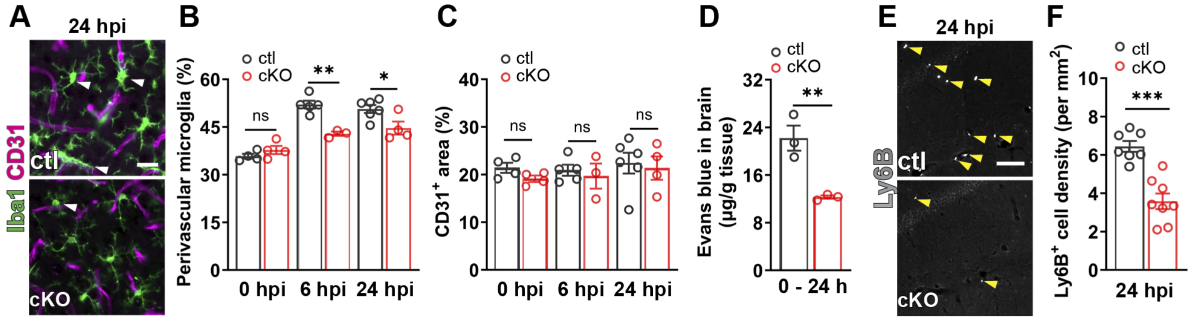


**Figure S5. Astrocytic A1AR deficiency reduces BBB disruption and neutrophil infiltration post peripheral LPS injection.**

**(A)** Representative images of immunolabeled Iba1$^+$ microglia and CD31$^+$ blood vessels at 24 hpi. Arrowheads indicate perivascular microglia. Scale bar = 20 µm.

**(B)** Proportion of perivascular microglia was increased in cKO mice compared to ctl mice (n = 4 mice in ctl/ cKO at 0 hpi, n = 5 mice in ctl at 6 hpi, n = 3 mice in cKO at 6 hpi, n = 6 mice in ctl at 24 hpi, n = 4 mice in cKO at 24 hpi).

**(C)** CD31$^+$ area was not altered in cKO and ctl mice at 0 hpi, 6 hpi, 24 hpi.

**(D)** EB extravasation was reduced in the brains of cKO mice compared to ctl mice which were injected with EB at 0 hpi and analyzed at 24 hpi (n = 3 mice per group).

**(E)** Representative images of immunolabeled of Ly6B$^+$ neutrophils in the brain parenchyma at 24 hpi. Scale bars = 20 µm.

**(F)** The density of Ly6B$^+$ cells was reduced in the brain of cKO mice compared to ctl mice at 24 hpi (n = 7 ctl mice and 8 cKO mice).

Summary data are presented as the mean ± SEM. Statistical significance in (**B**, **C**) were assessed using a two-way ANOVA. Statistical significance in(**D**, **F**) were assessed using unpaired Student's t test, ns: not significant, *P < 0.05, **P < 0.01, ***P < 0.001.

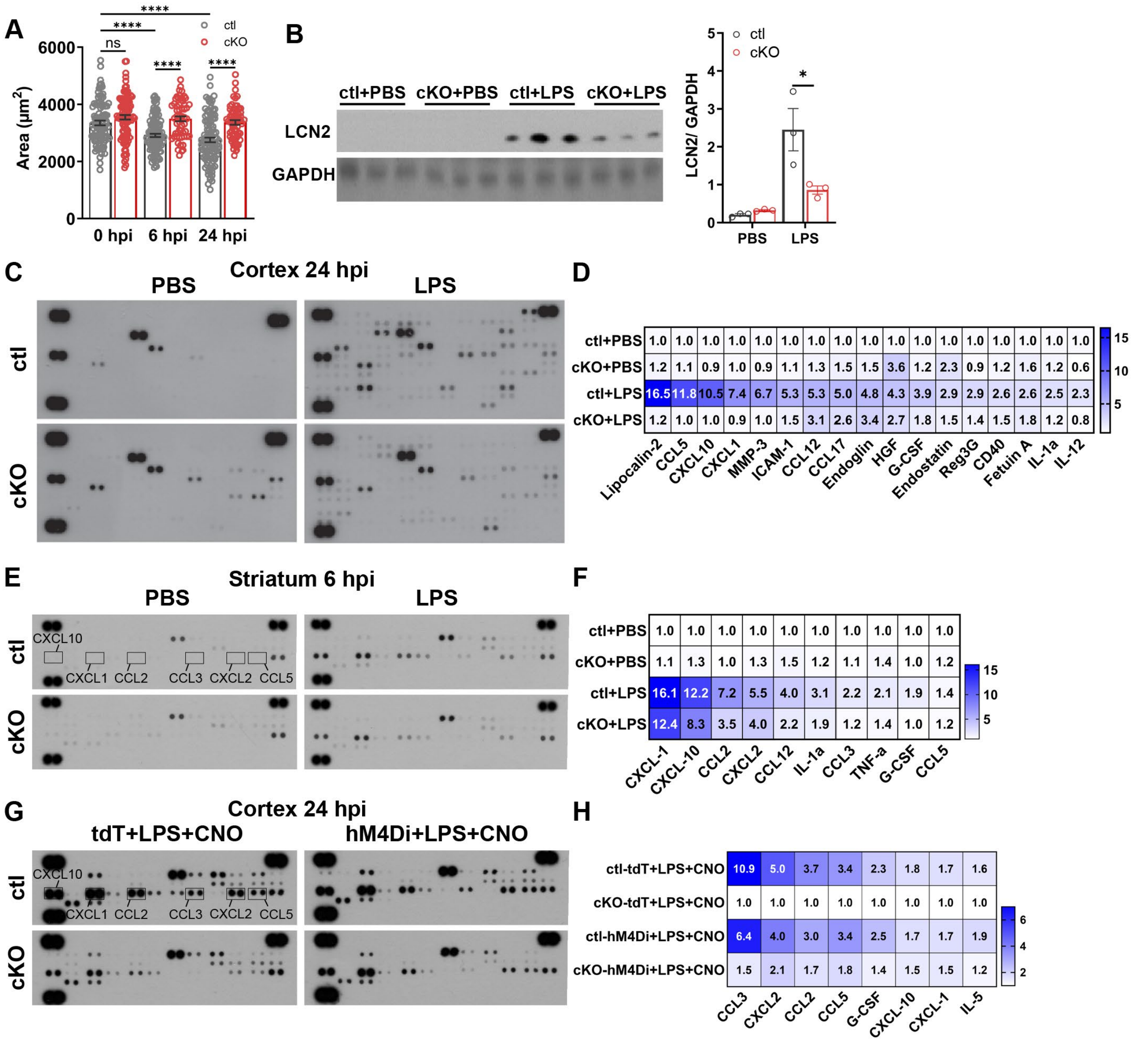


**Figure S6. Astrocytic A1AR deficiency reduces microglia reactivity and overall neuroinflammation upon LPS challenge which is attenuated by enhancing Gi signaling; related to Figure 4 and 7.**

**(A)** Microglial occupied area obtained from IMARIS-based morphological analysis of Iba1$^+$ microglia in cKO and ctl mice post LPS injection (n = 3 mice per group).

**(B)** LCN2 expression was reduced in cKO compared to ctl 24 hpi by Western blot.

**(C, D)** The expression of 111 cytokines in the cortex of ctl and cKO mice was measured by a proteomic profiling assay (**C**) at 24 h after PBS or LPS injection. Cytokine expression was reduced in the cortex of cKO group compared to ctl (PBS) group (**D**) (samples from 3 mice were mixed for each group).

**(E, F)** The expression of 40 cytokines in the striatum of ctl and cKO mice was measured by a proteomic profiling assay 6 hours after PBS or LPS injection (**E**). Cytokine expression was reduced in the cortex of cKO group compared to ctl (PBS) group (**F**) (samples from 3 mice were pooled for each group).

**(G, H)** The expression of 40 cytokines in the cortex of AAV-infected ctl and cKO mice was measured by a proteomic profiling assay 24 hours after LPS and CNO injection (**G**). Enhancing

Gi signaling in cKO mice increased cytokine expression after LPS and CNO injection (**H**) (samples from 3 mice were pooled for each group).
Summary data are presented as the mean ± SEM. Statistical significance in (**A, B**) were assessed by two-way ANOVA, ns: not significant, *P < 0.05, ****P < 0.0001.

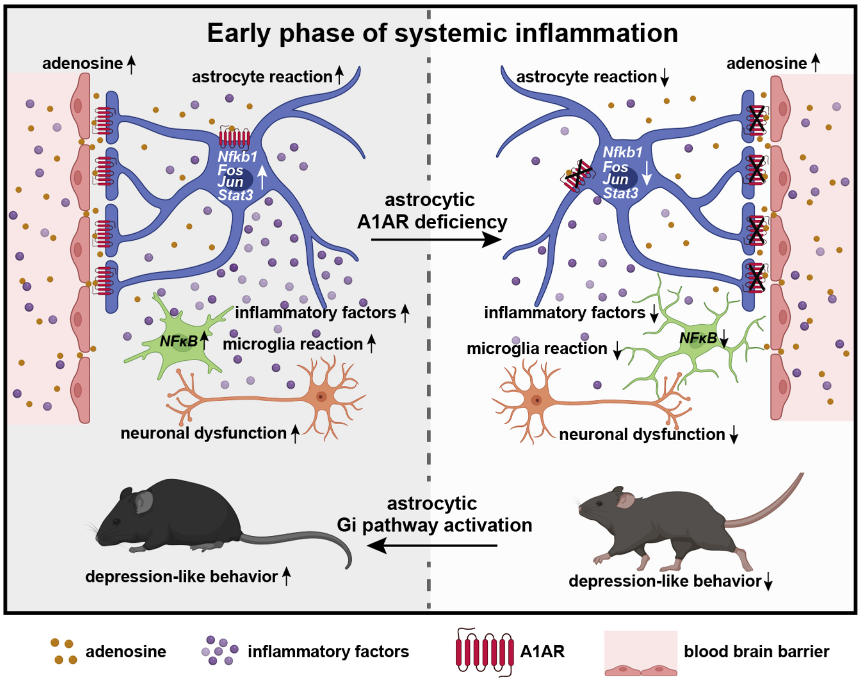
